# Zebrafish *mbnl* mutants model physical and molecular phenotypes of myotonic dystrophy

**DOI:** 10.1101/665380

**Authors:** Melissa N. Hinman, Jared I. Richardson, Rose A. Sockol, Eliza D Aronson, Sarah J. Stednitz, Katrina N. Murray, J. Andrew Berglund, Karen Guillemin

## Abstract

The muscleblind RNA binding proteins (MBNL1, MBNL2, and MBNL3) are highly conserved across vertebrates and are important regulators of RNA alternative splicing. Loss of MBNL protein function through sequestration by CUG or CCUG RNA repeats is largely responsible for the phenotypes of the human genetic disorder myotonic dystrophy (DM). We generated the first stable zebrafish (*Danio rerio*) models of DM-associated *MBNL* loss of function through mutation of the three zebrafish *mbnl* genes. In contrast to mouse models, zebrafish double and triple homozygous *mbnl* mutants were viable to adulthood. Zebrafish *mbnl* mutants displayed disease-relevant physical phenotypes including decreased body size and impaired movement. They also exhibited widespread alternative splicing changes, including the misregulation of many DM-relevant exons. Physical and molecular phenotypes were more severe in compound *mbnl* mutants than in single *mbnl* mutants, suggesting partially redundant functions of Mbnl proteins. The high fecundity and larval optical transparency of this complete series of zebrafish *mbnl* mutants will make them useful for studying DM-related phenotypes and how individual Mbnl proteins contribute to them, and for testing potential therapeutics.

## Introduction

The muscleblind (MBNL) family of RNA binding proteins (MBNL1, MBNL2, and MBNL3) is highly conserved in structure and function across multicellular species (Oddo *et al*, 2016). Vertebrate MBNL proteins contain two pairs of zinc finger motifs that bind to consensus YGCY sequences (Y = pyrimidine) in target RNAs and regulate multiple aspects of RNA metabolism, including alternative splicing (Ashwal-Fluss *et al*, 2014; Batra *et al*, 2014; Goers *et al*, 2010; Grammatikakis *et al*, 2011; Ho *et al*, 2004; Rau *et al*, 2011; Wang *et al*, 2012; Warf & Berglund, 2007). In general, MBNL proteins promote exon skipping or inclusion when bound to introns upstream or downstream of an alternative exon, respectively (Du *et al*, 2010; Goers *et al.*, 2010; Wang *et al.*, 2012). Regulation of alternative splicing by MBNL proteins influences the molecular and biological functions of hundreds of target genes.

Myotonic dystrophy types 1 and 2 (DM1 and DM2) are human genetic disorders caused primarily by MBNL protein loss of function. In DM, expression of expanded CTG or CCTG repeat RNAs leads to the formation of RNA stem loop structures, which contain dozens to thousands of YGCY motifs that sequester MBNL proteins, blocking their normal functions (Brook *et al*, 1992; Fardaei *et al*, 2002; Fu *et al*, 1992; Liquori *et al*, 2001; Mahadevan *et al*, 1992; Miller *et al*, 2000) (Fig 1A). DM1 disease severity increases with increasing CTG repeat length, likely due to increased MBNL protein sequestration (Yum *et al*, 2017). Although best known for its skeletal muscle phenotypes such as weakness, myotonia, atrophy, and pain, DM causes severe multi-systemic symptoms including cataracts, breathing difficulties, behavioral and psychological disorders, sleep disorders, insulin resistance, heart conduction abnormalities, cardiomyopathy, and alterations in the motility and microbiota of the gut (Bellini *et al*, 2006; Hilbert *et al*, 2017; Tarnopolsky *et al*, 2010; Thomas *et al*, 2018; Tieleman *et al*, 2008; Wenninger *et al*, 2018). Misregulation of specific alternative splicing events underlies disease phenotypes (Thomas *et al.*, 2018). For example, missplicing of *RYR1* and *ATP2A1* contributes to altered calcium homeostasis in DM1 muscle (Kimura *et al*, 2005; Zhao *et al*, 2015).

**Figure 1.**
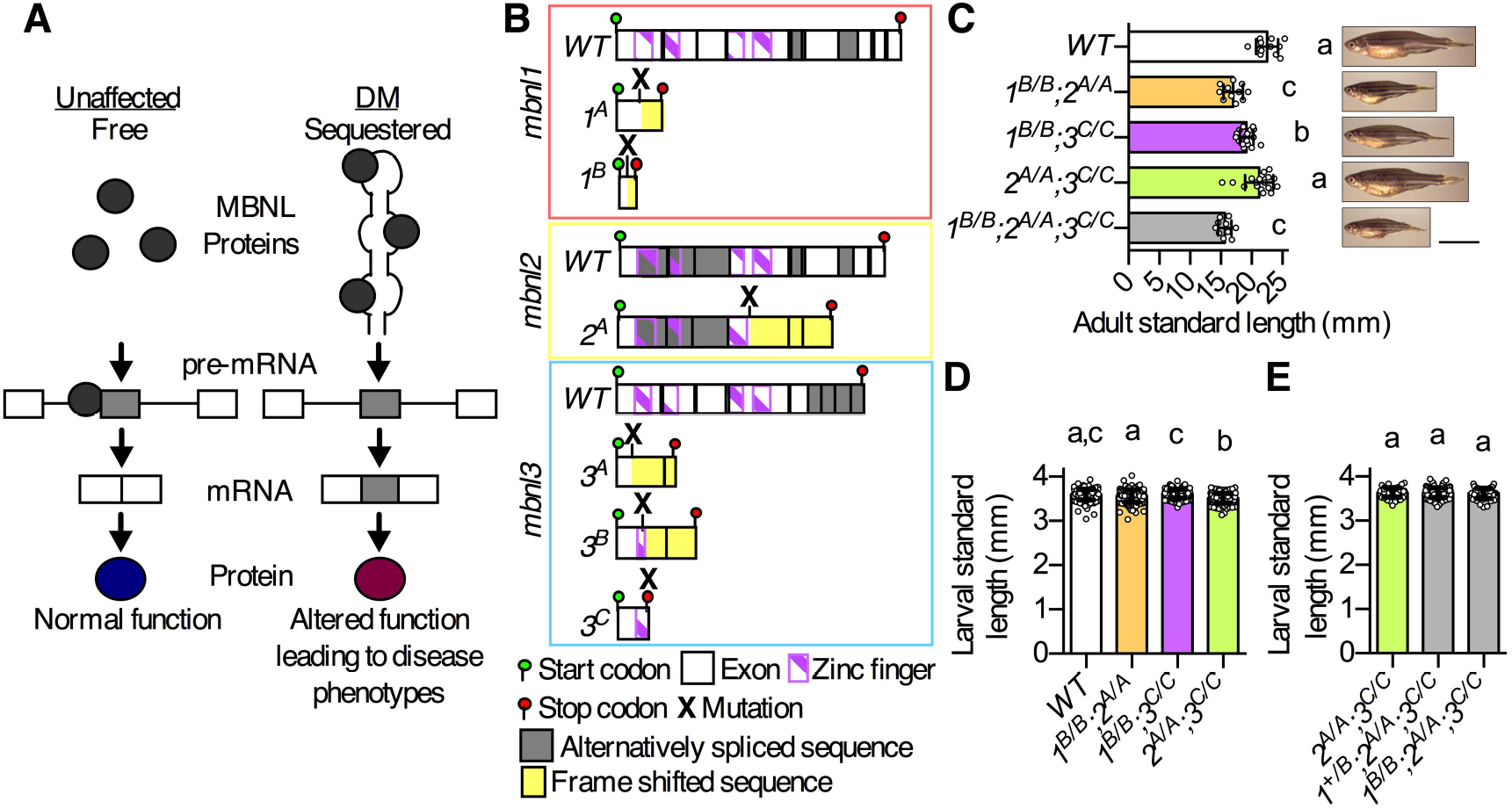
Mutation of zebrafish *mbnl* genes resulted in decreased adult body size. A. Primary molecular mechanism of the human genetic disorder myotonic dystrophy (DM). In unaffected individuals, free MBNL1, MBNL2, and MBNL3 proteins bind to target pre-mRNAs and regulate the inclusion of alternative exons in mRNAs. MBNL proteins suppress alternative exon inclusion in the example shown here, but they promote inclusion of other target alternative exons. In individuals with DM, MBNL proteins are sequestered by long CUG (DM1) or CCUG (DM2) repeat RNAs, which decreases their availability to bind to target pre-mRNAs and alters mRNA isoform production and downstream protein function. B. Diagram of WT and mutant zebrafish *mbnl1*, *mbnl2*, and *mbnl3* predicted coding sequences. *mbnl1* mutant alleles are denoted as *1^A^* and *1^B^*, *mbnl2* alleles as *2^A^*, and *mbnl3* alleles as *3^A^, 3^B^*, and *3^C^*. Mutant sequences are shown in Table EV2. C. Standard length of young adult 60 days post fertilization (dpf) WT and *mbnl* mutant zebrafish that were raised in the same tank. On the right are representative images of fish of each genotype taken at 76 dpf. Scale bar = 10 mm. D-E. Standard length of 7 dpf WT and (D) double *mbnl* mutant zebrafish or (E) clutchmates from an incross of *1^+/B^;2^A/A^;3^C/C^* fish. Data information: In (C-E), each dot represents one fish and data are presented as mean ± standard deviation. Data were analyzed by ordinary one-way ANOVA with Tukey’s multiple comparisons test. Data bars that do not share a letter above them are significantly different from one another.

*Mbnl* mutant mice have been invaluable for studying how individual Mbnl proteins contribute to DM-related phenotypes. Mouse *Mbnl1* is widely expressed, while *Mbnl2* expression is brain-enriched, and *Mbnl3* is expressed primarily during embryonic development and injury-induced adult skeletal muscle regeneration. *Mbnl1^−/−^* mice exhibit alternative splicing changes, myotonia, myopathy, and cataracts, but do not have other DM-related phenotypes such as muscle weakness (Kanadia *et al*, 2003; Lee *et al*, 2013). *Mbnl2^−/−^* mice exhibit DM-like CNS phenotypes including missplicing in the brain, abnormal sleep patterns, and spatial memory deficits (Charizanis *et al*, 2012). *Mbnl3^−/−^* mice exhibit delays in injury-induced muscle regeneration, accelerated onset of age-related pathologies, and gene expression changes, but display minimal changes in alternative splicing (Choi *et al*, 2016; Poulos *et al*, 2013). Compound *Mbnl* mutant mice have more severe phenotypes than single mutant mice, suggesting that Mbnl proteins have redundant functions. *Mbnl1^−/−^;Mbnl2^−/−^* mice are embryonic lethal, while *Mbnl1^−/−^;Mbnl2^+/−^* mice are viable but have severe missplicing, decreased body weight, myotonia, progressive muscle weakness, and reduced lifespan (Lee *et al.*, 2013). Muscle-specific double (*Mbnl1/Mbnl2*) and triple (*Mbnl1/Mbnl2/Mbnl3*) homozygous mutant mice are reduced in size, have dramatic and widespread alternative splicing changes, muscle weakness, and most die in the neonatal period due to respiratory distress (Thomas *et al*, 2017). Compound *Mbnl* mutant mice recapitulate many of the phenotypes associated with severe forms of DM, but their utility in experiments is limited due to difficulty in generating them in large numbers.

We used a complementary vertebrate organism, the zebrafish, to model DM-associated MBNL loss of function. Like mice, zebrafish have a single ortholog of each human *MBNL* gene (Liu *et al*, 2008). Unlike mice, zebrafish embryos are transparent and develop rapidly and externally, enabling direct studies of developmental phenotypes and of DM-related phenotypes such as altered gut motility and abnormal heart rhythm. In addition, zebrafish produce hundreds of embryos at once, and adults can be maintained more cheaply than mice, enabling experiments with large numbers of animals that improve the statistical power to study subtle or variable phenotypes. Two existing zebrafish DM models have severe limitations. Embryos that are transiently injected with CUG repeat RNA have subtle developmental abnormalities, but do not exhibit alternative splicing changes (Todd *et al*, 2014). Embryos in which *mbnl2* expression is temporarily knocked down using morpholinos, which often have off-target effects, exhibit profound morphological abnormalities that are inconsistent with the much milder phenotypes observed in *Mbnl2^−/−^* mice (Charizanis *et al.*, 2012; Machuca-Tzili *et al*, 2011). Neither zebrafish model can be used to study DM-relevant phenotypes beyond early development.

In this study, we made homozygous zebrafish *mbnl1*, *mbnl2*, and *mbnl3* loss of function mutants, which were crossed to generate double (*mbnl1/mbnl2*, *mbnl2/mbnl3*, and *mbnl1/mbnl3*) and triple (*mbnl1/mbnl2/mbnl3*) homozygous mutants, all of which were viable to adulthood. Zebrafish *mbnl* mutants exhibited DM-relevant physical phenotypes including decreased body size and impaired motor function. They also exhibited widespread alternative splicing changes, including many of the same changes that were present in DM patients and mouse *Mbnl* mutants. These alternative splicing changes occurred both in larval and adult fish, but were most dramatic in adult skeletal and heart muscle. As in mice, double and triple homozygous zebrafish *mbnl* mutants exhibited more severe phenotypes than single mutants. Thus, zebrafish *mbnl* mutants had physical and molecular phenotypes consistent with those present in DM, and are powerful new vertebrate models for studies of *MBNL* function.

## Results and Discussion

### A comprehensive set of zebrafish *mbnl* mutants was generated

To model DM-associated *MBNL* loss of function, we targeted constitutively included exons in each zebrafish *mbnl* gene using CRISPR, and established two *mbnl1* (*1^A/A^* and *1^B/B^*), one *mbnl2* (*2^A/A^*), and three *mbnl3* mutant lines (*3^A/A^*, *3^B/B^, 3^C/C^*) (Fig 1B, Tables EV1 and EV2) (Liu *et al.*, 2008). Although we lacked antibodies suitable for assessing zebrafish Mbnl protein levels (see materials and methods for panel of commercial antibodies tested), each *mbnl* mutation was predicted to result in loss of protein function due to frameshift and/or introduction of an early stop codon, leading to disruption of one or more of the zinc fingers that mediate RNA binding (Fig 1B and Table EV2) (Grammatikakis *et al.*, 2011). Homozygous larval mutants exhibited decreased (*2^A/A^*, *3^A/A^*, and *3^B/B^*) or unchanged (*1^A/A^*, *1^B/B^*, and *3^C/C^*) levels of the mRNA expressed from the mutated gene, and the expression of the other *mbnl* family members was largely unchanged (Fig EV1A-C), arguing against genetic compensation in these lines (El-Brolosy *et al*, 2019).

All homozygous *mbnl1, mbnl2*, and *mbnl3* mutants survived to adulthood in roughly Mendelian ratios (Appendix Table S1), were fertile, and did not exhibit the dramatic morphological phenotypes that were previously observed in *mbnl2* morpholino-injected larvae (Machuca-Tzili *et al.*, 2011). We propose that partially retained protein function due to the presence of one remaining zinc finger pair (Fig 1B) or morpholino off-target effects may explain the discrepancy in phenotype between *mbnl2* morphants and mutants. We favor the latter explanation, as Mbnl2 function was not required for survival in mice (Charizanis *et al.*, 2012). Double (*1^B/B^;2^A/A^, 1^B/B^;3^C/C^*, and *2^A/A^;3^C/C^*) and triple (*1^B/B^;2^A/A^;3^C/C^*) mutants were also viable, although triple mutants were present in lower than expected numbers and were unable to produce embryos except through *in vitro* fertilization (Appendix Table S1). This was, to our knowledge, the first time that *mbnl1/mbnl3* and *mbnl2/mbnl3* double homozygous mutant animals were generated. The viability of *1^B/B^;2^A/A^* and *1^B/B^;2^A/A^;3^C/C^* fish was surprising, as it contradicted findings in mouse models (Lee *et al.*, 2013; Thomas *et al.*, 2018). Most muscle-specific double and triple *Mbnl* mutant mice died during the neonatal period when they transitioned abruptly to breathing, but the small fraction that made it through this transition survived to adulthood (Thomas *et al.*, 2018). Perhaps *1^B/B^;2^A/A^* and *1^B/B^;2^A/A^;3^C/C^* fish were better able to survive because larval fish underwent a gradual transition between receiving oxygen through diffusion and through the gills and/or because the presence of partially functional Mbnl2 protein was sufficient for survival. In summary, we created the first complete panel of vertebrate *mbnl1, mbnl2*, and *mbnl3* single, double, and triple homozygous mutants for modeling DM.

### Mutation of *mbnl* genes led to decreased zebrafish size

We asked whether zebrafish had similar phenotypes to mice, in which compound, but not single, *Mbnl* mutants were significantly smaller than WT (Lee *et al.*, 2013; Thomas *et al.*, 2017). We raised WT, double, and triple homozygous *mbnl* mutant fish under identical conditions and measured their lengths in early adulthood. WT fish were significantly larger than *1^B/B^;3^C/C^* fish, which in turn were larger than *1^B/B^;2^A/A^* and *1^B/B^;2^A/A^;3^C/C^* fish (Fig 1C). Although *2^A/A^;3^C/C^* fish were slightly smaller than WT fish, the difference was not statistically significant (Fig 1C). Mutation of *mbnl1*, *mbnl2*, or *mbnl3* alone was not sufficient to decrease adult zebrafish size (Fig EV2A-C). Taken together, these results suggested that zebrafish Mbnl proteins have partially redundant functions, as the growth defect was more dramatic in compound mutants than in single mutants.

We also measured 7 days post fertilization (dpf) larval fish prior to exogenous feeding. All double homozygous *mbnl* mutant larvae were similar in size to WT, except for *2^A/A^;3^C/C^*, which were slightly smaller than WT (Fig 1D). *1^B/B^;2^A/A^;3^C/C^* larvae were similar in size to their *1^+/B^;2^A/A^;3^C/C^* and *2^A/A^;3^C/C^* clutch mates, and all single homozygous larvae were either similar in size or slightly larger than WT (Fig 1E and Fig EV2D). These results suggest that the reduced size phenotype in zebrafish *mbnl* mutants arose later in development.

Histological analysis of adult skeletal muscle that was performed by a fish pathologist indicated that the fish genotypes with the most profound size phenotypes, *1^B/B;^2^A/A^* and *1^B/B^;2^A/A^;3^C/C^*, did not exhibit obvious signs of the myofiber atrophy or centralized nuclei that were described in compound *mbnl* mutant mice and in individuals with DM (Fig EV3) (Thomas *et al.*, 2018; Thomas *et al.*, 2017). We cannot, however, rule out the possibility that the mutant fish had subtle alterations in muscle structure or function.

### Zebrafish *mbnl* mutants had altered movement

Given that DM impairs motor function, we examined whether zebrafish *mbnl* mutants exhibited altered swimming behavior by introducing individual adult fish to a novel tank and tracking their position over five minutes (Fig 2A and B). An analogous test was performed in DM patients, in which the distance walked in six minutes was used as a measure of motor function (Kierkegaard & Tollback, 2007; Park *et al*, 2018). While all single homozygous mutants swam equal or greater distance than WT, two of the double homozygous *mbnl* mutants (*1^B/B^;2^A/A^* and *1^B/B^;3^C/^*^C^) and the *1^B/B^;2^A/A^;3^C/C^* mutants swam significantly decreased distances compared to WT (Fig 2C-E). One possible explanation for this result is impaired motor function, perhaps due to subtle changes in muscle structure and function that were not apparent in histological analysis.

**Figure 2.**
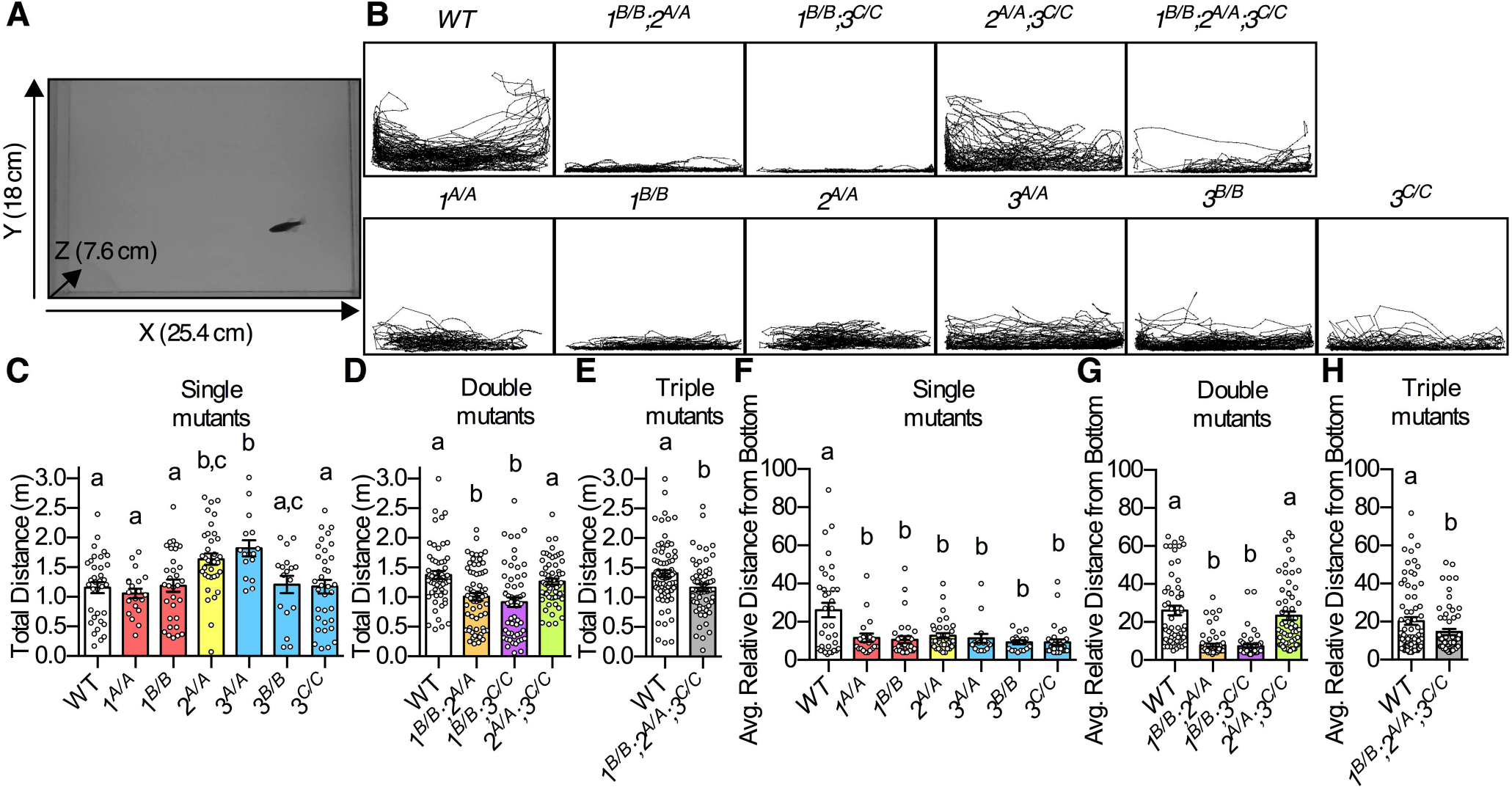
Zebrafish *mbnl* mutants exhibited altered movement. A. Example still image taken from video of swim tests. Software was used to track the center of mass of each fish in the X and Y directions over 5 minutes. B. Representative traces of individual fish of each genotype taken during the swim tests. X and Y axes are the same as in (A). *mbnl1* mutant alleles are denoted as *1^A^* and *1^B^*, *mbnl2* alleles as *2^A^*, and *mbnl3* alleles as *3^A^, 3^B^*, and *3^C^*. C-E. Total distance that adult WT and (C) single, (D) double, and (E) triple homozygous *mbnl* mutant fish swam during a five-minute swim test. F-H. Average relative distance from the bottom of the tank of fish during a five-minute swim test. Zero represents the bottom of the tank and 100 represents the top of the tank. Data information: In (C-H) each dot represents one fish and data are presented as mean ± SEM. In (C,D,F,G) data were analyzed by ordinary one-way ANOVA with Tukey’s multiple comparisons test and in (E,H) data were analyzed by a Student’s t-test. Data bars that do not share a letter above them are significantly different from one another.

We also observed a striking difference in the positions of *mbnl* mutants and WT fish within the tank, which was quantified by measuring the distance from the bottom of the tank in each frame for each fish and averaging this over the recording period. Except for *2^A/A^;3^C/C^* mutants, all *mbnl* mutants spent significantly more time toward the bottom of the tank than WT fish (Fig 2F-H). Avoiding the top of the tank is a well-known indicator of anxiety in zebrafish, and future tests will explore whether *mbnl* mutants exhibit other signs of anxiety or perhaps have a morphological or motor deficit that explains their lower swimming position (Cachat *et al*, 2010). Intriguingly, individuals with DM often show signs of avoidant personality disorder with anxiety, and perhaps the *mbnl* mutant fish model that phenotype (Delaporte, 1998; Minier *et al*, 2018). Overall, these results indicated that loss of *mbnl* function resulted in altered movement in fish, but the exact mechanisms still need to be explored. Impaired feeding due to altered movement is a potential explanation for the size phenotype observed in young adult *mbnl* mutant fish.

### Zebrafish *mbnl* mutants displayed changes in alternative splicing across tissues

Given the DM-relevant size and movement phenotypes of *mbnl* mutant fish, we asked whether they also exhibited DM-associated alternative splicing changes. The model system used in our initial studies was the extensively characterized suppression of *mbnl1* and *mbnl2* exon 5 inclusion by Mbnl proteins, which is known to affect the subcellular localization and splicing regulatory activity of the encoded proteins (Gates *et al*, 2011; Terenzi & Ladd, 2010; Tran *et al*, 2011). The intronic sequences immediately upstream of *mbnl1* and *mbnl2* exon 5, which contain putative YGCY Mbnl protein binding sites, are highly conserved between human and zebrafish, suggesting functional importance (Appendix Figs S1 and S2).

We harvested total RNA from whole larval zebrafish and adult skeletal muscle, heart, brain, cornea, and intestine. Quantitative RT-PCR analyses of WT fish indicated that *mbnl1, mbnl2*, and *mbnl3* mRNAs were present in larvae and across adult tissues, and that levels of all three were highest in skeletal muscle and brain (Fig EV1D-F). Inclusion of *mbnl1* exon 5 and *mbnl2* exon 5, as measured by RT-PCR, was significantly elevated in zebrafish *mbnl* mutants across all tissues (Figs 3 and EV4). The phenotype was most dramatic in skeletal muscle and heart, where *mbnl1* and *mbnl2* exon 5 were almost entirely skipped in WT and predominantly included in *1^B/B^;2^A/A^* and *1^B/B^;2^A/A^;3^C/C^* mutants (Figs 3B and C and EV4B and C). The brain and cornea exhibited *mbnl* splicing patterns similar to one another, but with lower magnitudes changes than those observed in muscle (Figs 3D and E and EV4D and E). The corneal splicing phenotype suggests that zebrafish *mbnl* mutants may also model the genetic disorder Fuchs endothelial corneal dystrophy, a subtype of which was recently shown to be caused by an expanded CUG repeat that is associated with MBNL protein sequestration and mis-splicing (Figs 3 and EV4)(Mootha *et al*, 2017; Mootha *et al*, 2015; Wieben *et al*, 2017; Winkler *et al*, 2018). Modest but significant splicing changes were observed in whole larvae and intestine (Figs 3A and F and EV4A and F).

**Figure 3.**
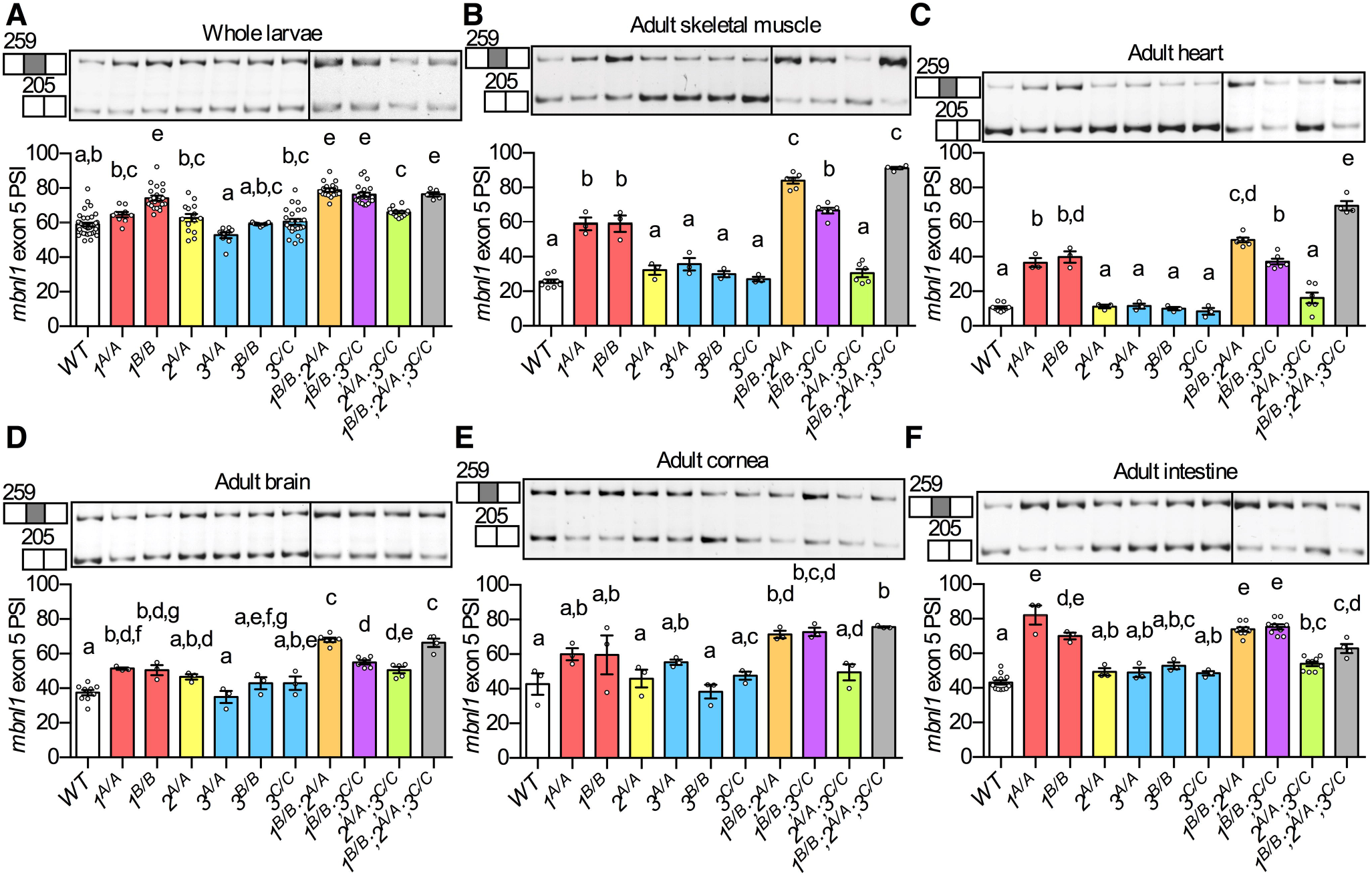
Alternative splicing of *mbnl1* exon 5 was misregulated across tissues in zebrafish *mbnl* mutants. A-F. RT-PCR analysis showing percent spliced in (PSI) of *mbnl1* exon 5 in WT and *mbnl* mutant (A) whole 5 dpf larvae and in adult (B) skeletal muscle, (C) heart, (D) brain, (E) cornea, and (F) intestine. *mbnl1* mutant alleles are denoted as *1^A^* and *1^B^*, *mbnl2* alleles as *2^A^*, and *mbnl3* alleles as *3^A^, 3^B^*, and *3^C^*. Representative RT-PCR gels are shown above each graph with band sizes in bp on the left. Dividing lines indicate samples run on separate gels. Data information: In (A-F) each dot represents RNA from one adult fish or a pool of five larval fish. Data are presented as mean ± SEM. Data were analyzed by ordinary one-way ANOVA with Tukey’s multiple comparisons test. Data bars that do not share a letter above them are significantly different from one another.

In most tissues, double and triple mutants exhibited larger *mbnl1* and *mbnl2* splicing changes than single mutants (Figs 3 and EV4). This is consistent with the idea that zebrafish Mbnl proteins, like mouse Mbnl proteins, have partially redundant functions when it comes to splicing regulation (Lee *et al.*, 2013; Thomas *et al.*, 2017). Strikingly, the genotypes with the strongest splicing phenotypes, *1^B/B^;2^A/A^* and *1^B/B^;2^A/A^;3^C/C^*, also had dramatic size and movement phenotypes (Figs 1-3 and EV4).

In mouse model and DM patient tissues, some regulated exons were much more sensitive than others to changes in overall levels of free MBNL proteins (Wagner *et al*, 2016). Zebrafish also followed this pattern. For example, in skeletal muscle and heart, *mbnl1* mutation alone was sufficient to increase *mbnl1* exon 5 inclusion, whereas mutation of both *mbnl1* and *mbnl2* was required to increase *mbnl2* exon 5 inclusion (Figs 3B and C and EV4B and C). Overall, these results suggest that, as in other systems, zebrafish Mbnl proteins work in concert to regulate alternative splicing across tissues.

### Misregulation of alternative splicing was widespread in zebrafish *mbnl* mutants

To understand the genome-wide impact of *mbnl* mutation on alternative splicing, we performed RNA-Seq analysis using RNA isolated from the skeletal muscle of adult WT, *1^B/B^, 2^A/A^, 3^C/C^, 1^B/B^;2^A/A^, 1^B/B;^3^C/C^, 2^A/A^;3^C/C^*, and *1^B/B^;2^A/A^;3^C/C^* zebrafish. There were no significant changes in the levels of total *mbnl1*, *mbnl2*, or *mbnl3* mRNAs in mutants compared to WT (Fig EV1G-I). Hundreds of significantly misregulated RNA alternative splicing events were detected in all *mbnl* mutants, with misregulated cassette exons outnumbering alternative 3’ and 5’ splice sites, retained introns, and mutually exclusive exons (Fig 4A and Table EV3). The widespread splicing changes in all mutants suggested that all three zebrafish *mbnl* proteins contributed to splicing regulation, and that each *mbnl* mutation caused at minimum a partial loss of function. *1^B/B^;2^A/A^* and *1^B/B^;2^A/A^;3^C/C^* fish had the largest number of significantly misregulated splicing events (Fig 4A). Surprisingly, *2^A/A^;3^C/C^* mutants exhibited fewer overall dysregulated alternative splicing events than *2^A/A^* mutants or *3^C/C^* mutants (Fig 4A). In mouse models, many cassette exon events were shown to be dysregulated in opposite directions in *Mbnl3* mutants and other *Mbnl* mutants (Thomas *et al.*, 2017). Perhaps opposing splicing regulation could be a contributing factor to the apparent mild splicing and tank position phenotypes of *2^A/A^;3^C/C^* mutants.

**Figure 4.**
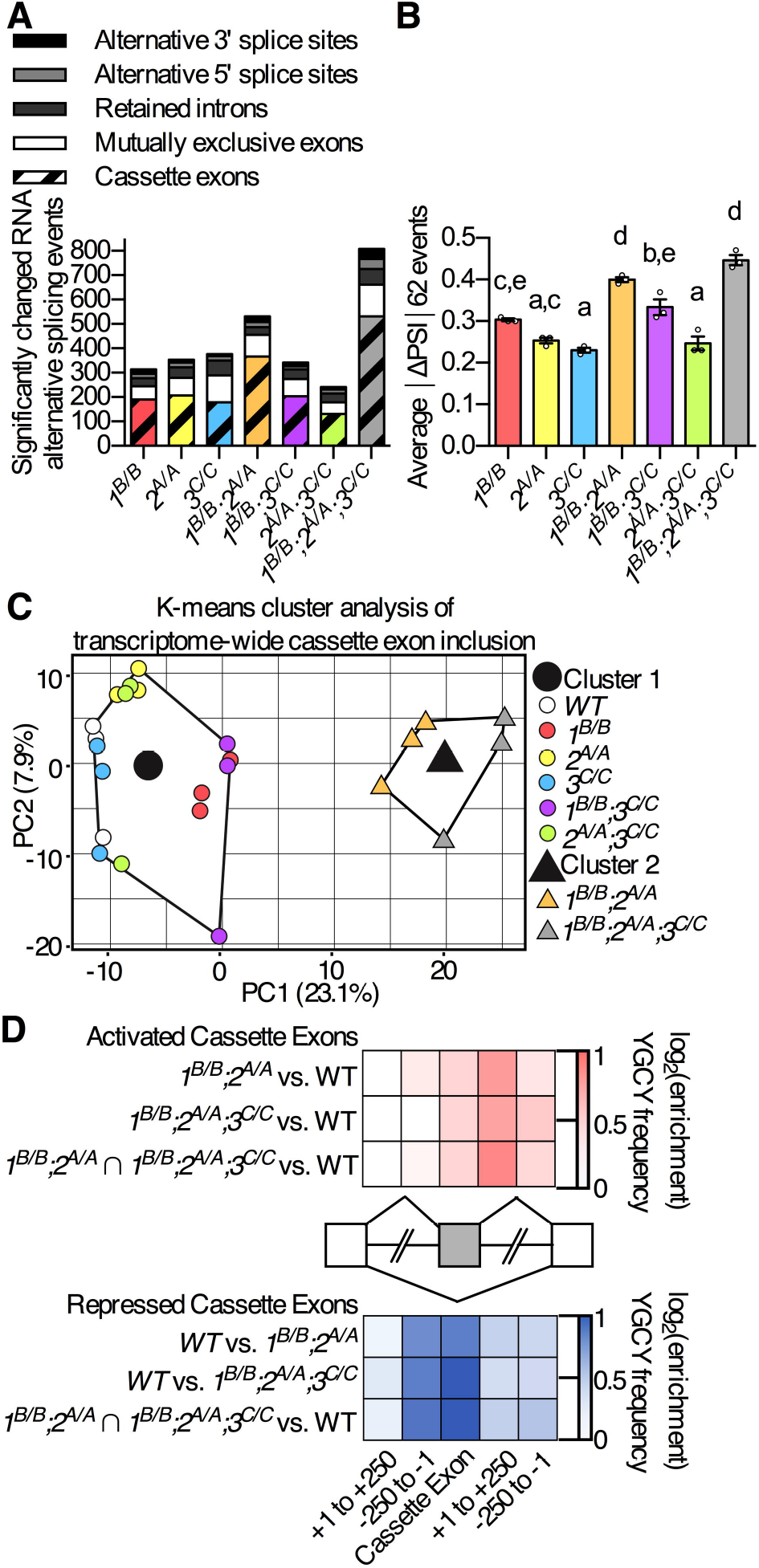
Zebrafish *mbnl* mutations led to widespread changes in adult skeletal muscle RNA alternative splicing. A. Total number of RNA alternative splicing events of different types that were significantly misregulated between WT and *mbnl* mutant adult skeletal muscle, as identified by RNA-Seq. B. The average absolute value of the change in percent spliced in (ΔPSI) is shown for the set of 62 cassette exons whose inclusion was significantly misregulated in RNA-Seq analysis of the skeletal muscle of at least four of seven *mbnl* mutant fish lines compared to WT. C. K-means cluster analysis based on changes in cassette exon inclusion showing that *1^B/B^;2^A/A^* and *1^B/B^;2^A/A^;3^C/C^* mutants cluster closely with each other, while other mutants cluster more closely with WT. Each small circle or triangle represents an individual fish, and the large circle and triangle represent the centers of the clusters. D. Heat maps showing the enrichment of the previously identified Mbnl protein binding sequence YGCY (where Y is a pyrimidine) within significantly misregulated cassette exons, in the intronic sequences 250 bp immediately upstream and downstream of those exons, and in the 250 bp of intronic sequences immediately adjacent to the flanking constitutive exons. Activated cassette exons are those in which Mbnl proteins regulated inclusion positively, and repressed cassette exons are those for which inclusion is decreased by Mbnl proteins. The YGCY enrichment analysis was performed for the set of cassette exons that were significantly misregulated in *1^B/B^;2^A/A^* fish, in *1^B/B^;2^A/A^;3^C/C^* fish, and for the set of misregulated exons that overlapped between the two (*1^B/B^;2^A/A^* ⍰*1^B/B^;2^A/A^;3^C/C^*). Data information: In (A-D) the *mbnl1* mutant allele is denoted as*1^B^*, the *mbnl2* allele as *2^A^*, and the *mbnl3* allele as *3^C^*. In (B) each dot represents one fish. Data are presented as mean ± SEM. Data were analyzed by ordinary one-way ANOVA with Tukey’s multiple comparisons test. Data bars that do not share a letter above them are significantly different from one another.

We also analyzed the magnitude of the splicing change (absolute value of the change in percent spliced in (ΔPSI) between WT and mutant) for the 62 cassette exon events that were significantly misregulated in the majority of *mbnl* mutants (at least 4 of 7). The highest magnitude of splicing change across events was also observed in *1^B/B^;2^A/A^* and *1^B/B^;2^A/A^;3^C/C^* fish (Fig 4B). These results suggest that zebrafish Mbnl proteins, like mouse orthologs, have at least partially redundant functions (Lee *et al.*, 2013; Thomas *et al.*, 2017).

We next asked which genotypes were most similar in their overall splicing phenotypes. All cassette exon splicing changes that were observed in any mutant compared to WT were compiled into a single list, and the top 800 events with the largest variability in PSI were identified. Using the gap statistic, we determined that all samples could be clustered into two groups based on the splicing phenotype for those events. A K-means cluster analysis with two centers was then performed, in which all WT and mutant samples were analyzed for their similarity in PSI for the 800 events. The analysis indicated that all *1^B/B^;2^A/A^* and *1^B/B^;2^A/A^;3^C/C^* fish were more similar to each other than they were to WT fish or any of the other *mbnl* mutant fish (Fig 4C). Taken together, these results indicated that compound loss of function of zebrafish Mbnl proteins led to widespread and robust changes in alternative splicing.

We analyzed the frequency of the Mbnl protein binding motif, YGCY, within and surrounding the cassette exons that were significantly misregulated in *1^B/B^;2^A/A^* and *1^B/B^;2^A/A^;3^C/C^* mutants as well as those misregulated cassette exons that were in common between the two mutants. YGCY motifs were enriched in the introns downstream of exons whose inclusion was activated by the presence of Mbnl proteins, while YGCY motifs were enriched upstream and within the cassette exons that were repressed by Mbnl proteins (Fig 4D). This was consistent with findings in other DM model systems, and suggests that zebrafish Mbnl proteins played a direct role in regulating the inclusion of cassette exons (Du *et al.*, 2010; Goers *et al.*, 2010; Oddo *et al.*, 2016; Wang *et al.*, 2012).

### Zebrafish *mbnl* mutants exhibited disease-relevant alternative splicing changes

Given the widespread changes in alternative splicing in *mbnl* mutant zebrafish, we asked whether these changes were conserved with those identified in human DM patients. Using publicly available datasets, we identified cassette exons whose inclusion was significantly misregulated in DM1 tibialis anterior muscle biopsy tissues or in DM1 patient derived myoblasts compared to healthy control tissues, and compared them with the zebrafish RNA-Seq data. We identified 25 orthologous cassette exons that were misregulated in both *1^B/B^;2^A/A^;3^C/C^* mutant fish and in DM1 tibialis muscle and 40 that were misregulated in both *1^B/B^;2^A/A^;3^C/C^* mutant fish and in the DM1 myoblasts (Fig 5A and EV5A and Source Data Fig 5A and EV5A). The inclusion of these alternative exons tended to be misregulated in the same direction in zebrafish DM models and in DM1 patient-derived tissues (Figs 5A and EV5A).

**Figure 5.**
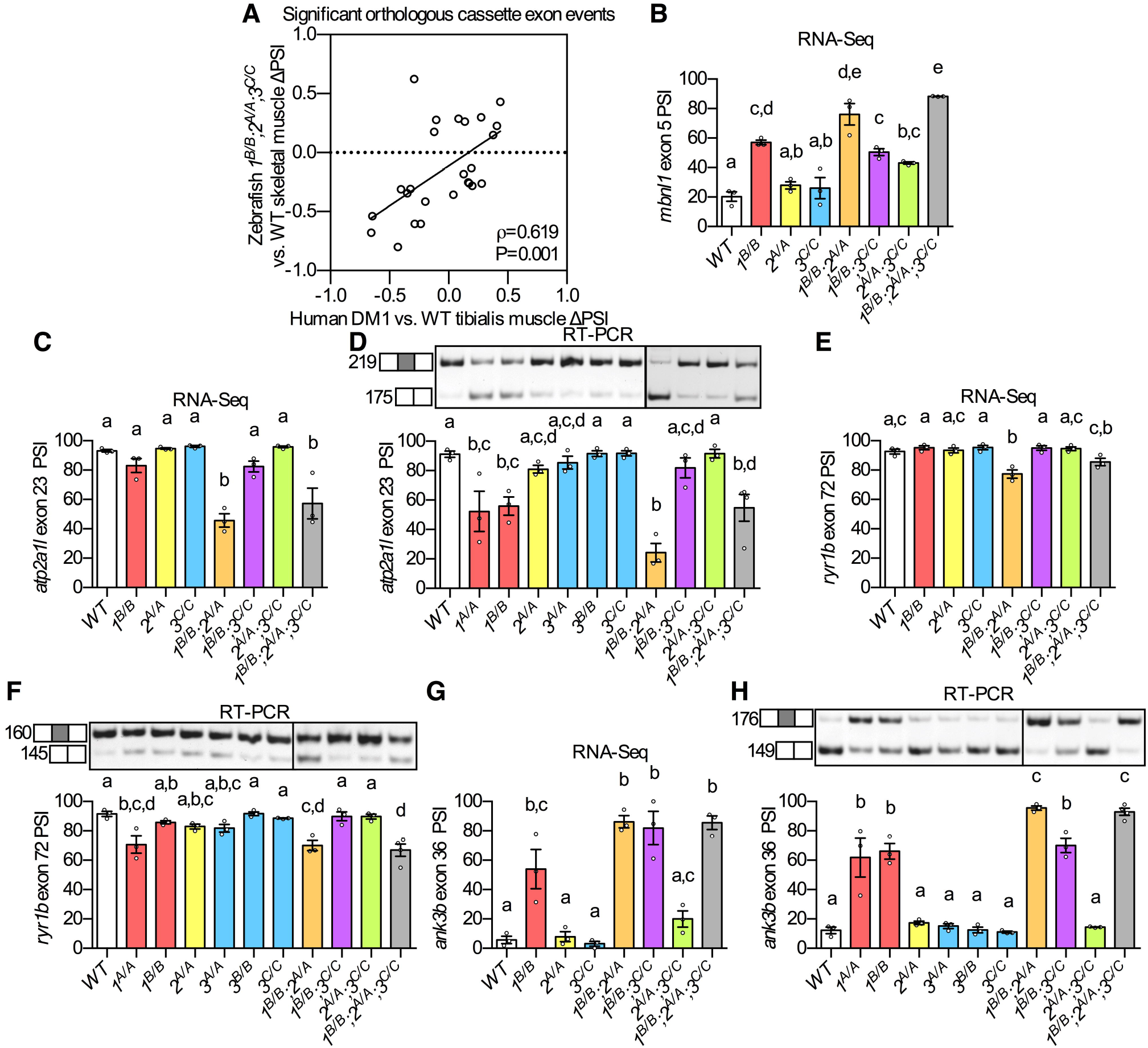
Misregulation of many DM-associated alternative splicing events was conserved in *mbnl* mutants. A. Change in percent spliced in values (ΔPSI) between mutant and WT are shown for orthologous exons in zebrafish *1^B/B^;2^A/A^;3^C/C^* skeletal muscle and in tibialis muscle from human DM1 patients. B-H. (B,C,E,G) RNA-Seq and (D,F,H) RT-PCR analyses showing percent spliced in (PSI) of (B) *mbnl1* exon 5, (C-D) *atp2a1l* exon 23, (E-F) *ryr1b* exon 72, and (G-H) *ank3b* exon 36 in WT and *mbnl* mutant adult zebrafish skeletal muscle. There is published evidence of misregulation of the human orthologs of each of these exons in DM patient-derived tissues (Freyermuth *et al.*, 2016; Kimura *et al.*, 2005; Zhao *et al.*, 2015). Data information: In (A-H) *mbnl1* mutant alleles are denoted as *1^A^* and *1^B^*, *mbnl2* alleles as *2^A^*, and *mbnl3* alleles as *3^A^, 3^B^*, and *3^C^*. In (A) ρ is the Spearman’s rank correlation coefficient. In (B-H) data are presented as mean ± SEM. Each dot represents RNA from one fish. Representative gels are shown above each RT-PCR graph with band sizes in bp shown on the left. Dividing lines indicate samples run on separate gels. Data were analyzed by ordinary one-way ANOVA with Tukey’s multiple comparisons test. Data bars that do not share a letter above them are significantly different from one another.

As anticipated, *mbnl1* exon 5 and *mbnl2* exon 5 appeared on both lists of orthologous misregulated cassette exons, and PSI values were strikingly similar as determined by RNA-Seq and RT-PCR (Figs 3B, 5B, EV4B, and EV5B). Two other randomly selected orthologous exons, *aplp2* exon 7 and *atp6v1h* exon6, also showed similar splicing phenotypes by RNA-Seq and RT-PCR, and contained potential YGCY Mbnl binding motifs in their surrounding intronic sequences (Fig EV5C-F and Appendix Figs S3 and S4). Figs 5A and EV5A do not represent an exhaustive list of orthologous misregulated exons, as a manual literature search revealed several additional orthologs of disease-associated exons that were misregulated in *mbnl* mutant model fish, some of which were only annotated in an earlier zebrafish genome assembly (GRCz10, Table EV3). For example, the decreased inclusion of alternative exons within the human *ATP2A1* and *RYR1* genes contributes to altered calcium homeostasis in DM muscle, and we detected decreased inclusion of orthologous exons in zebrafish *mbnl* mutants (Fig 5C-F, and Appendix Figs S5 and S6)(Kimura *et al.*, 2005; Zhao *et al.*, 2015). Likewise, we observed dramatic mis-splicing of zebrafish *ank3b* exon 36, the human ortholog of which is known to be misregulated in human DM1 heart samples (Fig 5G and H and Appendix Fig S7)(Freyermuth *et al*, 2016). Taken together, these results indicate that the overall coordinated splicing regulation program of the Mbnl proteins was well-conserved between zebrafish and humans, although not every individual missplicing event was conserved. For example, *mbnl* mutants did not exhibit changes in alternative splicing of the zebrafish orthologs of *CLCN1*, whose splicing misregulation contributes to myotonia in DM. Thus, the zebrafish may not be an appropriate model for studying myotonia, although it can be used to study many other important disease phenotypes (Lueck *et al*, 2007a; Lueck *et al*, 2007b).

### Zebrafish *mbnl* mutants are powerful models for studying DM

For the first time, we have modeled DM-associated *MBNL* loss of function by generating a complete panel of vertebrate *mbnl* single, double, and triple homozygous mutants. These fish exhibited molecular and physical phenotypes similar to those observed in humans with DM and mouse models, including decreased body size, impaired movement, and widespread changes in alternative splicing. We propose that zebrafish *1^B/B^;2^A/A^* and *1^B/B^;2^A/A^;3^F/F^* mutants, which exhibited the most dramatic phenotypes, represent severe forms of DM1 with long CTG repeats in which most, but probably not all, MBNL proteins are sequestered, while other double and single *mbnl* mutants model more moderate forms of the disease in which MBNL sequestration is less robust.

These new zebrafish models, and the accompanying RNA-Seq data that we generated, will be valuable in future studies of how individual Mbnl proteins and specific alternative splicing changes contribute to DM-relevant phenotypes. For example, exons that were misregulated in all zebrafish *mbnl* mutants except for *2^A/A^;3^C/C^* mutants are strong candidates for contributing to the anxiety-like phenotype (Fig 2F-H). Likewise, exons whose mis-regulation was not conserved between humans with DM and zebrafish are candidates for contributing to disease-related changes in muscle histology that were not observed in the zebrafish *mbnl* mutants (Fig EV3). Zebrafish DM models complement other existing vertebrate model systems because hundreds of larval or adult fish can be generated easily to study subtle or variable phenotypes and to test potential therapeutics. Additionally, transparent larval zebrafish can be used to directly study disease-related phenotypes, such as altered gut motility and heart abnormalities, in live animals.

## Materials and methods

### Generation of mutant zebrafish

All zebrafish experiments were performed with the guidance and approval of the University of Oregon Institutional Animal Care and Use Committee (PHS assurance number D16-00004, protocol AUP-15-98). Guide RNAs (gRNAs) targeting zebrafish (*Danio rerio*) *mbnl1*, *mbnl2*, and *mbnl3* were designed using the Chop Chop website (http://chopchop.cbu.uib.no). DNA templates for the gRNAs were generated by a template-free Phusion polymerase (New England Biolabs) PCR reaction using a common scaffold primer (gRNA scaffold, Table EV1) and a gene-specific primer (*mbnl1* gRNA1, *mbnl1* gRNA2, *mbnl2* gRNA, *mbnl3* gRNA1, or *mbnl3* gRNA2, Table EV1), then cleaned using the QIAquick PCR Purification Kit (Qiagen). gRNAs were transcribed from DNA templates using a MEGAscript kit (Ambion) and purified by phenol-chloroform extraction and isopropanol precipitation. Cas9 RNA was generated by linearizing the pT3TS-nls-zCas9-nls plasmid (Jao *et al*, 2013) with XbaI, purifying it using the QIAquick Gel Extraction Kit (Qiagen), performing an in vitro transcription reaction using the T3 mMESSAGE kit (Invitrogen), and purifying the RNA using the RNeasy Mini kit (Qiagen). AB zebrafish embryos were microinjected at the one cell stage with 1-2 nL of a mixture containing 100 ng/μL Cas9 mRNA, 50 ng/μL gRNA, and phenol red, and raised to adulthood. Mosaic mutants were identified by PCR amplification and Sanger sequencing of fin clip DNA using primers specific to the targeted region (Tables EV1 and EV2), and outcrossed to wildtype AB zebrafish to generate heterozygotes. Fish with predicted loss of function mutations were identified by Sanger sequencing (Table EV2), and further crossed to generate single, double, and triple *mbnl1*, *mbnl2*, and *mbnl3* homozygotes. Fish were genotyped by restriction fragment length polymorphism analysis using the primers and restriction enzymes indicated in Table EV2. Sperm from all zebrafish mutant lines were cryopreserved and are available upon request from the corresponding author.

### Measurement of zebrafish size

Young adult fish (2-4 months old) (Fig 1C and Fig EV2A-C) were anesthetized in 168 mg/L tricaine methane sulfonate, photographed against a white background with a ruler using a tablet computer, finclipped, and genotyped. To ensure identical density and feeding conditions, fish were compared with others from the same clutch (Fig EV2A-C) or different clutches that had been raised together in the same tank (Fig 1C). For measurement of larval fish, unfed embryos of different genotypes were raised to 7 days post fertilization (dpf) in separate dishes at a density of one fish per mL of embryo medium, with the exception of the larvae in Fig 1E, which were grown in the same dish and genotyped after measurement. Fish were anesthetized in 168 mg/mL tricaine methane sulfonate, laid out on a microscope slide in 3% methylcellulose, and imaged using a Leica M165FC microscope. An investigator that was blinded to genotype measured the distance from snout to caudal peduncle or posterior tip of the notochord (standard length, SL) of each fish (Parichy *et al*, 2009). Data were analyzed by ordinary one-way ANOVA with multiple comparison correction (Source Data Figs 1 and EV2).

### Alternative splicing analysis by RT-PCR

Fish were euthanized by tricaine methane sulfonate overdose (larvae) or hypothermic shock (adults). Tissues of interest were dissected and flash frozen in 1 mL of Trizol (Ambion), thawed, and homogenized with a mortar and pestle (larvae) or Bullet Blender Storm 24 (adult tissues). Chloroform (200 μL) was added to each tube followed by mixing, centrifugation at 12,000 g for 10 minutes at 4°C, transfer of the aqueous phase to a separate tube, addition of 200 μL ethanol, and binding of sample to an RNeasy mini kit column (Qiagen). RNA was washed and eluted according to the manufacturer’s instructions and the concentration was measured using a NanoDrop 2000 (Thermo Scientific).

RNA (20-200 ng) was reverse transcribed with Superscript II Reverse Transcriptase (Invitrogen) according to the manufacturer’s instructions using gene-specific reverse (R) primers located in the exon downstream of the regulated exon of interest (Table EV1). The resulting cDNA was amplified by 28-31 cycles of PCR using Taq polymerase and the forward (F) and reverse (R) primers indicated in Table EV1. Samples were separated by electrophoresis on a 6% *bis*-Acrylamide (19:1) gel that was stained overnight with 1X SYBR green I nucleic acid gel stain (Invitrogen). The gel was imaged and quantified using an AlphaImagerHP (Alpha Innotech). The background-corrected sum of each band was measured and the percent exon inclusion was calculated using the following formula: ((exon included sum) /(exon included sum + exon excluded sum)) × 100. Data were analyzed by ordinary one-way ANOVA with multiple comparison correction.

### RNA-Seq analysis

RNA was extracted using the RiboPure RNA Purification Kit (Invitrogen AM1924) from epaxial skeletal muscle from the tails of three biological replicates of adult WT and mutant fish. RNA quality was examined using the Fragment Analyzer RNA Analysis DNF-471 kit (Advanced Analytical) and all RQN values were > 8.0. Ribosomal RNA was depleted from 200 μg of the RNA using the NEBNext rRNA Depletion Kit (NEB E6310X) and then a cDNA library was prepared using the NEBNext Ultra RNA Library Prep Kit for Illumina (NEB E7530L). The libraries were checked for quality using the Fragment Analyzer NGS Analysis DNF-474 kit (Advanced Analytical) and quantitated using the KAPA Library Quantification Kit (KAPA Code KK4824). The completed libraries were then pooled in equimolar amounts and sequenced on an Illumina Next-Seq 500. A minimum of 60 million paired-end 75 × 75 reads were obtained for each library.

BCL files were demultiplexed and converted to fastq files using BCL2Fastq (version 2.16.0.10). The fastq files were then checked for quality using FastQC (version 0.11.8) and aligned to the GRCz11 zebrafish genome using STAR (version 2.5.1b). The alternative splicing events were then analyzed using rMATS (version 4.0.2), grouping the biological replicates of each mutant and comparing them against WT. A similar preliminary analysis of the same sequencing data was performed using GRCz10 and rMATS (version 4.0.2). Significant mis-splicing events were categorized as having a false discovery rate (FDR) < 0.10. Cluster analysis was performed with R and visualized with the factoextra package (version 1.0.5) using the top 800 most variable cassette exon splicing events in PSI values. Data were first tested using the gap statistic and then K-Means clustering was performed with two centers. The data for human DM1 tibialis anterior comparisons were downloaded from NCBI GEO, accession # GSE86356. Data for the DM1-derived myoblast cell line were downloaded from NCBI SRA, study # SRP158284. The data were processed as described above and then compared to the zebrafish data for orthologous mis-splicing events. Orthologous exons were found using a custom python script and using tblastx to confirm. Exons were counted as orthologous if the genes from which they were transcribed were at least 75% conserved between species, the exon was in the same place in the transcript (equal exon number in at least one annotated transcript), and if the exon had an evalue from tblastx of lower than 0.05.

Enrichment of putative YGCY Mbnl protein binding motifs was analyzed for the sets of cassette exons whose inclusion was significantly increased (FDR < 0.05, ΔPSI > 0.2) or decreased (FDR < 0.05, ΔPSI < −0.2) compared to WT in *1^B/B^;2^A/A^* mutants, *1^B/B^;2^A/A^;3^C/C^* mutants, or in both (*1^B/B^;2^A/A^* ∩ *1^B/B^;2^A/A^;3^C/C^*). For each set of cassette exons, the frequency of YGCY motifs (GCTT, CGCT, TGCT, and GCGC) was compared to the frequency of control 4-mers with identical A+T and CpG content within the following regions: cassette exons (normalized for analyzed sequence length), intronic sequences 1-250 bp immediately 5’ or 3’ of the cassette exons, intronic sequences 1-250 bp 3’ of the upstream constitutive exons, and intronic sequences 1-250 bp 5’ of the downstream constitutive exons. The log_2_ ((enrichment of YGCY motifs relative to control motifs in each region associated with regulated cassette exons)/(enrichment of YGCY motifs relative to control motifs in each region associated with non-regulated exons)) was plotted in heatmap form.

### Analysis of mbnl gene expression

RT-qPCR was used to measure mbnl RNA levels in zebrafish tissues. Total RNA was prepared from 5 dpf larvae and adult tissues using the same procedure as for alternative splicing analysis. RNA was treated with TURBO DNase (Ambion) according to manufacturer’s instructions then reverse transcribed with an oligo(dT)_20_ primer using the Superscript III cDNA First Strand Synthesis kit (Invitrogen). The qPCR reaction was set up using the KAPA SYBR FAST ABI Prism kit (KAPA Biosystems) according to manufacturer’s instructions and run on a Quant Studio 3 System (ThermoFisher) using the default settings for SYBR Green reagents and the fast run mode in the QuantStudio Design & Analysis Software v.1.4.2. The comparative C_T_ (ΔΔC_T_) method was used to calculate relative mRNA levels. Five biological samples of each genotype or tissue type were run in triplicate and data were normalized to the expression of the housekeeping gene *eef1a1l1*(Vanhauwaert *et al*, 2014). Primers used for RT-qPCR are shown in Table EV1.

Western blots analyses were performed using 15-50 μg of protein lysates that were prepared in RIPA buffer (Boston Bioproducts) with complete mini EDTA-free protease inhibitor (Roche) from HeLa cells (positive control for MBNL1 and MBNL2 protein expression), whole 5 dpf zebrafish larvae, adult skeletal muscle, heart, brain, intestine, and cornea, or purchased from Santa Cruz Biotechnology (human placenta extract used as a positive control for MBNL3 protein expression). The antibodies tested included Santa Cruz anti-MBNL1(D-4) sc-515374 (1:250), Santa Cruz anti-MBNL1 (4A8) sc-136165 (1:500), Millipore anti-MBNL1 ABE-241 (1:1000), Abcam anti-MBNL2 ab105331 (1:250), Sigma-Aldrich anti-MBNL3 SAB1411751 (1:150), and Sigma-Aldrich anti-actin A5060 (1:1000). Santa Cruz goat anti-rabbit IgG-HRP sc-2004 (1:2000) and Sigma goat anti-mouse IgG FC specific HRP (1:2000) were used as secondary antibodies.

### Zebrafish behavior assays

Age-matched single adult zebrafish were placed into a novel environment consisting of a custom glass aquarium measuring 18 cm deep × 25.4 cm long × 7.6 cm wide (Fig 2A). The fish were monitored with a Logitech camera during a 5-minute time frame to characterize basic exploratory swimming behavior. The raw tracking data was analyzed with custom software (DaniOPEN, https://github.com/stednitzs/daniopen) to measure swim distance and distance from the bottom of the tank (Stednitz *et al*, 2018). Following the test period, the fish were returned to their home tanks.

### Histology

Ten-month-old WT and mutant adult zebrafish were euthanized by hypothermic shock and fixed in Bouin’s solution (Sigma), then washed in 70% ethanol. Fixed fish were bisected parasagitally or transversely, processed for paraffin embedding, sectioned at 7 μm thickness, and stained with hematoxylin and eosin by a histologist from the University of Oregon Institute of Neuroscience. Muscle morphology was analyzed by a fish pathologist from the Zebrafish International Resource Center who was blinded to genotype. Representative DIC images of muscle transverse sections were taken using a Leica DMLB microscope.

## Data availability

RNA-Seq data are available on NCBI Gene Expression Omnibus accession number GSE145270, https://www.ncbi.nlm.nih.gov/geo/query/acc.cgi?acc=GSE145270.

## Acknowledgements

We thank the staff of the University of Oregon Zebrafish Facility for maintaining fish and providing expertise in performing zebrafish procedures, and Dr. Poh Kheng Loi for histological services. We thank Ellie Melancon, Camila Morales Fenero, and Michelle Massaquoi for advice in performing experiments and data analysis, and members of the labs of Karen Guillemin, Andy Berglund, and Judith Eisen for helpful discussions. Research reported in this publication was supported by the National Institute of General Medical Sciences of the National Institutes of Health (https://www.nigms.nih.gov) under award numbers 1P50GM098911 and 1P01GM125576 to K.G. and 5R01GM121862 to J.A.B., and by the National Institute of Diabetes and Digestive and Kidney Diseases of the National Institutes of Health (https://www.niddk.nih.gov) under award number F32DK107318 to M.N.H. It was also supported by grants from the Muscular Dystrophy Association (www.mda.org) (516314 to J.A.B. and 627218 to M.N.H.) and the Myotonic Dystrophy/Wyck Foundation (www.myotonic.org) (Wyck-FF-2014-0013 to M.N.H.). The content is solely the responsibility of the authors and does not necessarily represent the official views of the National Institutes of Health, the Muscular Dystrophy Association, or the Myotonic Dystrophy/Wyck Foundation. The funders had no role in study design, data collection and analysis, decision to publish, or preparation of the manuscript.

## Author contributions

M.N.H. – conceptualization, methodology, formal analysis, investigation, writing-original draft preparation, visualization, funding acquisition

J.R. – investigation, formal analysis, methodology, writing-original draft preparation

R.A.S. – investigation E.DA. – investigation

S.J.S. – investigation, resources

K.N.M. – investigation

J.A.B. – conceptualization, resources, writing-review and editing, supervision, project administration, funding acquisition

K.G. – conceptualization, resources, writing-review and editing, supervision, project administration, funding acquisition

## Expanded view figure legends

**Figure EV1.**
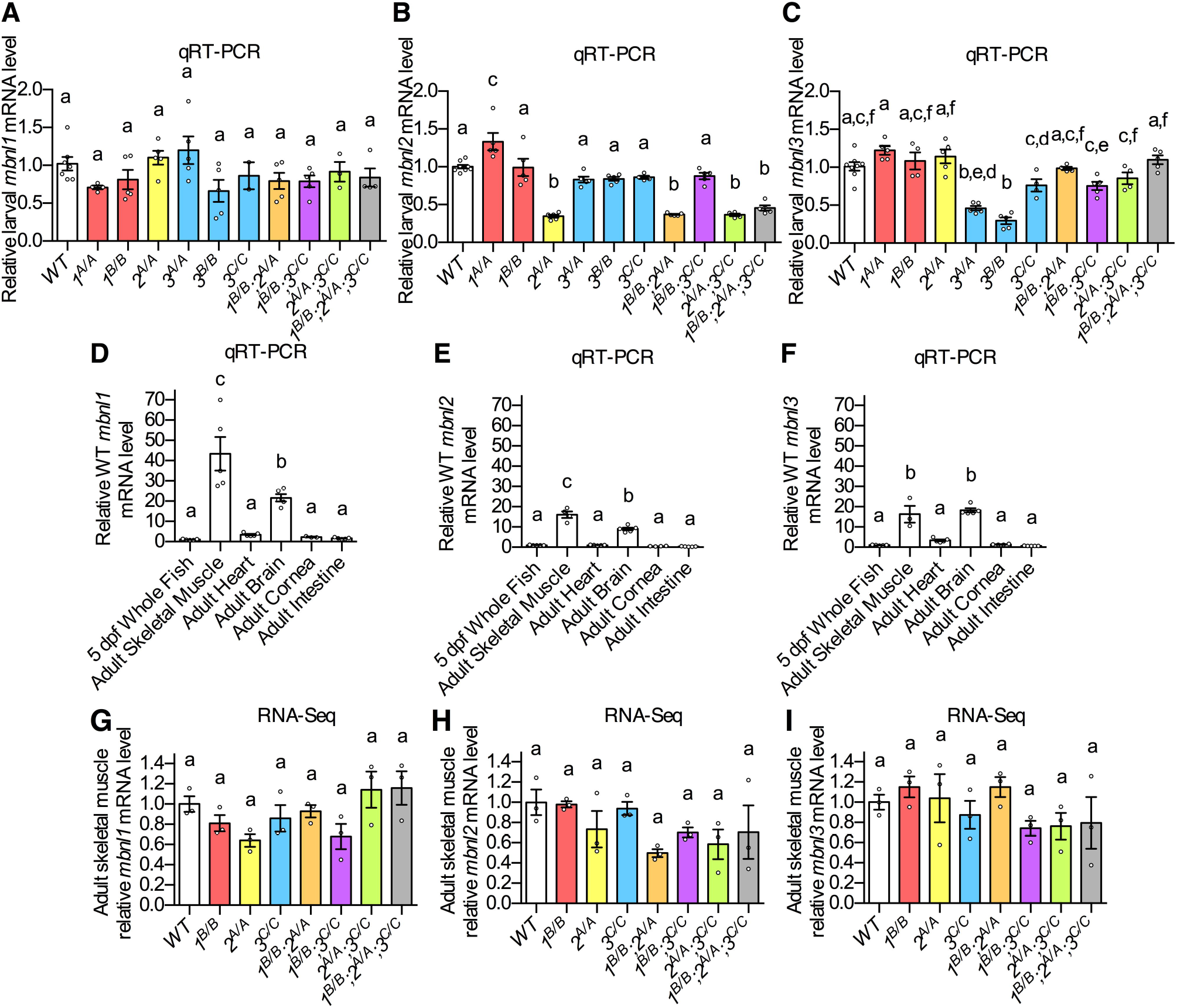
Effects of *mbnl* mutations on *mbnl* mRNA levels. A-C. qRT-PCR showing the levels of (A) *mbnl1*, (B) *mbnl2*, and (C) *mbnl3* mRNAs in WT and *mbnl* mutant 5 days post fertilization whole larval zebrafish. D-F. qRT-PCR showing relative levels of (D) *mbnl1* mRNA, (E) *mbnl2* mRNA, and (F) *mbnl3* mRNA in WT 5 dpf whole larvae, adult skeletal muscle, heart, brain, cornea, and intestine. G-I. RNA-Seq data showing the levels of (G) *mbnl1*, (H) *mbnl2*, and (I) *mbnl3* mRNAs in WT and *mbnl* mutant adult zebrafish skeletal muscle. Data information: In (A-I) data are presented as mean ± SEM. Each dot represents RNA from a pool of 5 larvae or one adult fish. Data were analyzed by ordinary one-way ANOVA with Tukey’s multiple comparisons test. Data bars that do not share a letter above them are significantly different from one another.

**Figure EV2.**
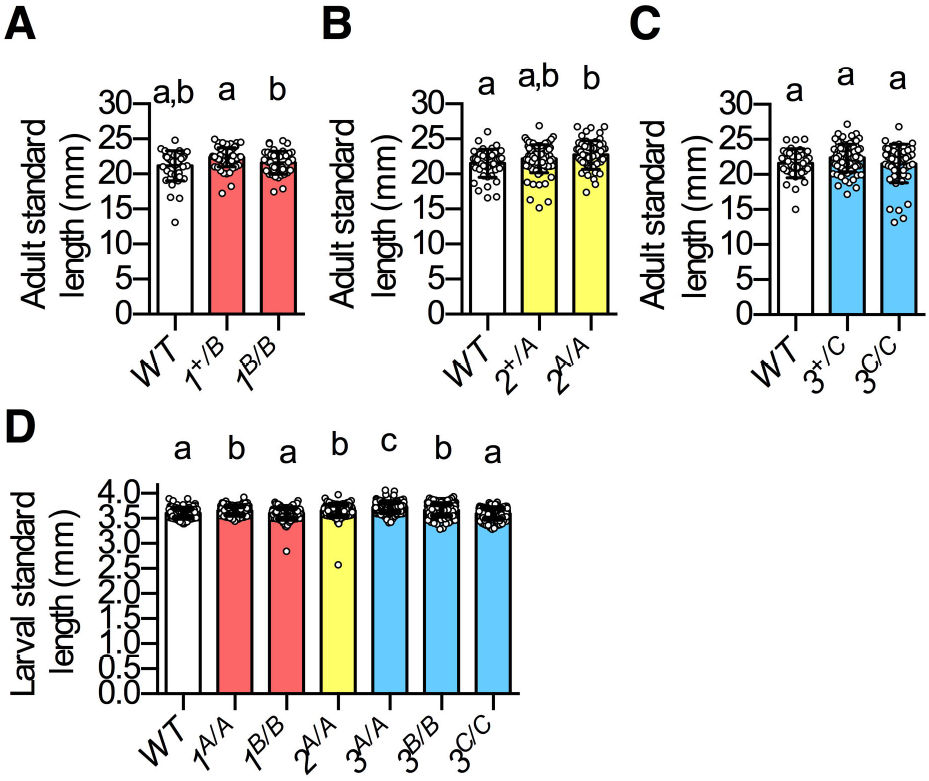
Single *mbnl* mutant zebrafish were not decreased in size. A-C. Standard length of (A) 83 dpf clutchmates from an incross of *1^+/B^* fish, (B) 83 dpf clutchmates from an incross of *2^+/A^* fish, and (C) 78 dpf clutchmates from an incross of *3^+/C^* fish. D. Standard length of 7 dpf larval fish. Data information: In (A-D) *mbnl1* mutant alleles are denoted as *1^A^* and *1^B^*, *mbnl2* alleles as *2^A^*, and *mbnl3* alleles as *3^A^, 3^B^*, and *3^C^*. Data are presented as mean ± standard deviation. Each dot represents one fish. Data were analyzed by ordinary one-way ANOVA with Tukey’s multiple comparisons test. Data bars that do not share a letter above them are significantly different from one another.

**Figure EV3.**
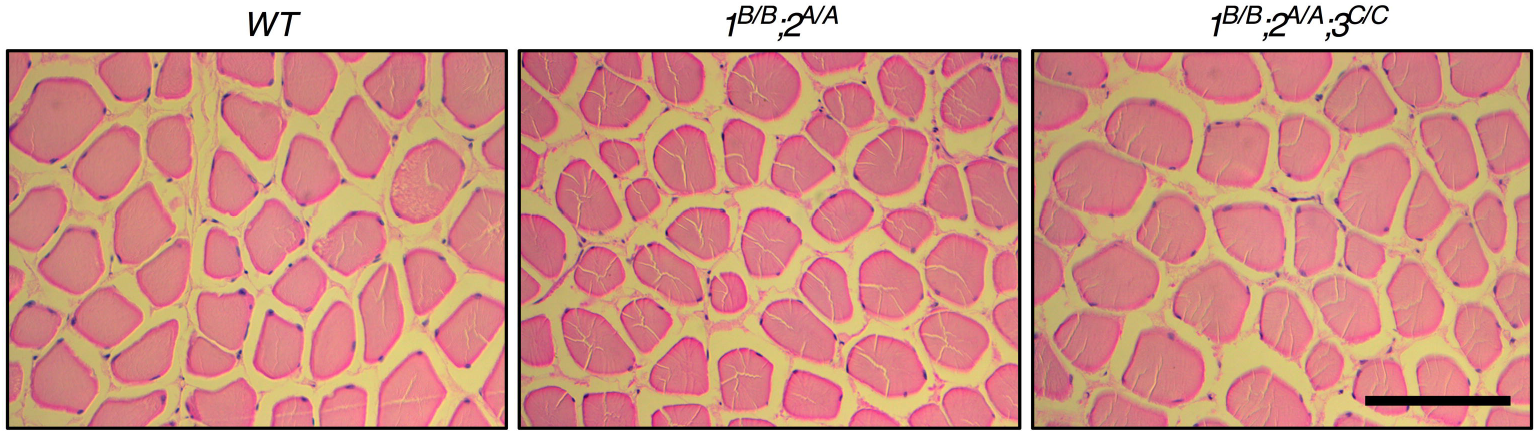
Adult zebrafish WT and *mbnl* mutant muscle fibers were structurally similar. Representative hematoxylin and eosin stained transverse sections of epaxial muscle fibers from the tails of adult zebrafish. Scale bar = 100 μm.

**Figure EV4.**
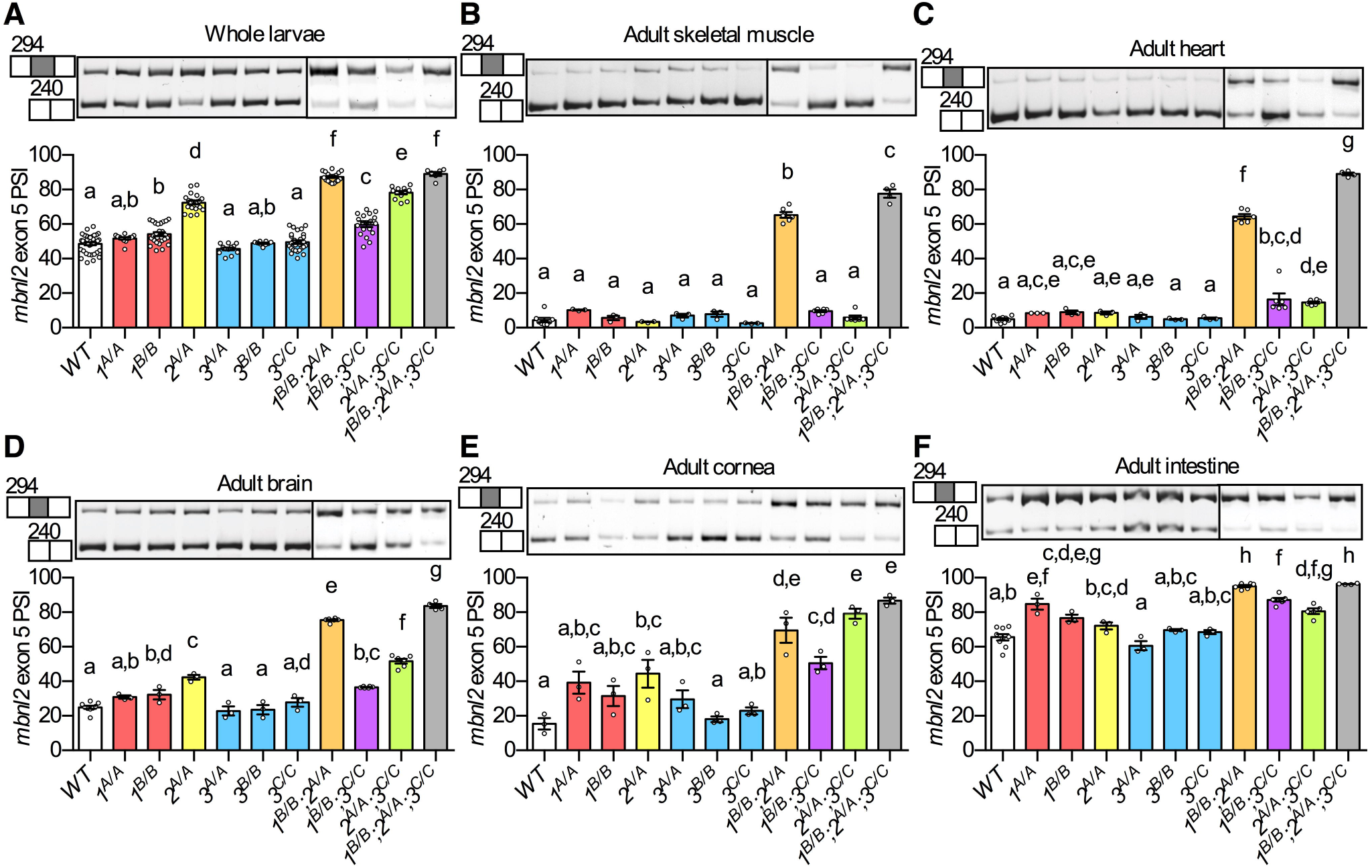
Alternative splicing of *mbnl2* exon 5 was misregulated across tissues in zebrafish *mbnl* mutants. A-F. RT-PCR analysis showing percent spliced in (PSI) of *mbnl2* exon 5 in WT and *mbnl* mutant (A) whole 5 dpf larvae, and in adult (B) skeletal muscle, (C) heart, (D) brain, (E) cornea, and (F) intestine. Data information: In (A-F) *mbnl1* mutant alleles are denoted as *1^A^* and *1^B^*, *mbnl2* alleles as *2^A^*, and *mbnl3* alleles as *3^A^, 3^B^*, and *3^C^*. Representative RT-PCR gels are shown above each graph with band sizes in bp shown on the left. Dividing lines indicate samples run on separate gels. Each dot represents RNA from one adult fish or a pool of five larvae. Data are presented as mean ± SEM. Data were analyzed by ordinary one-way ANOVA with Tukey’s multiple comparisons test. Data bars that do not share a letter above them are significantly different from one another.

**Figure EV5.**
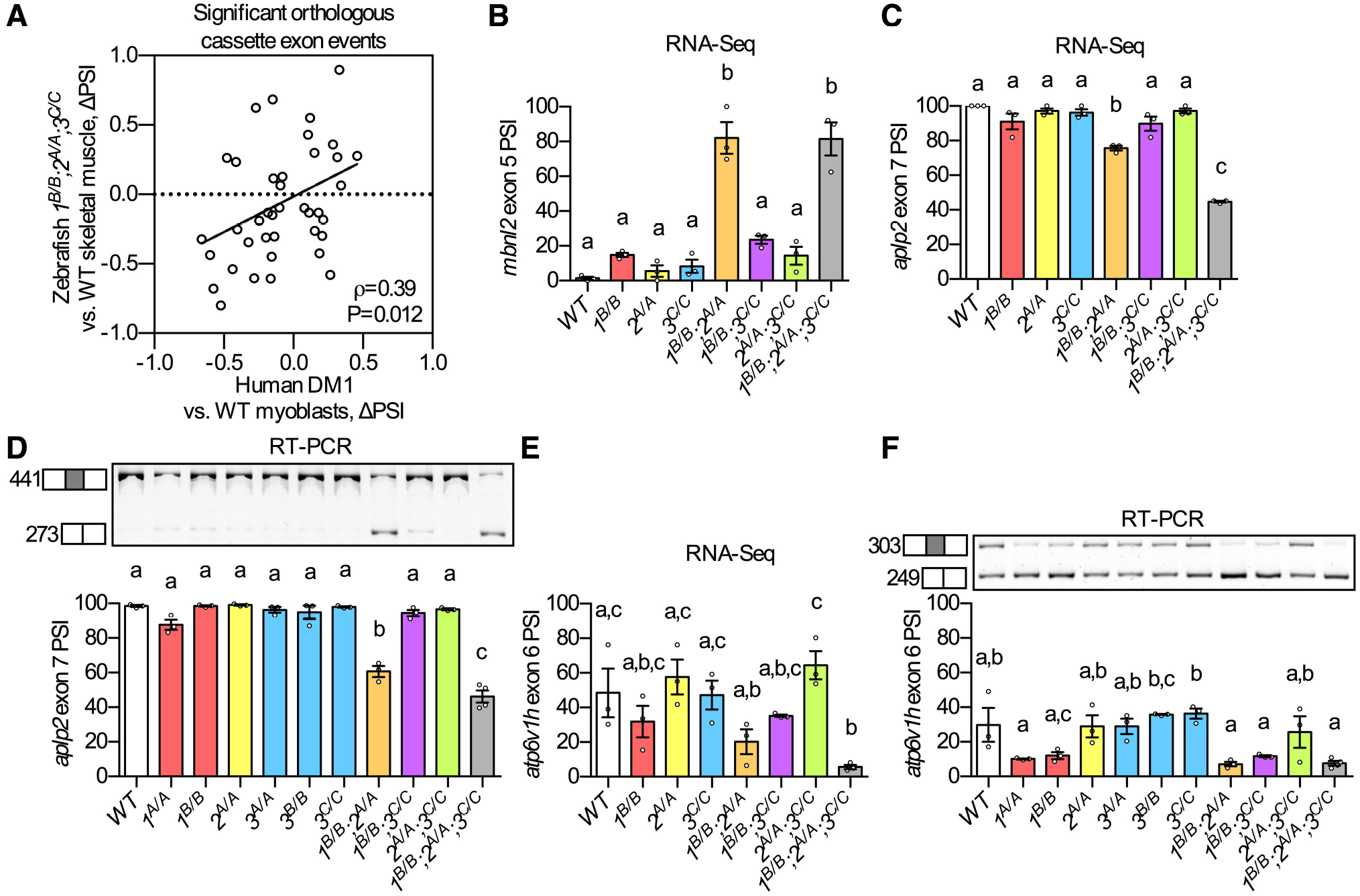
Misregulation of many DM-associated alternative splicing events was conserved in zebrafish *mbnl* mutants. A. Change in percent spliced in values (ΔPSI) between mutant and WT are shown for orthologous exons in zebrafish *1^B/B^;2^A/A^;3^C/C^* skeletal muscle and in human DM1 patient-derived myoblasts. B-F. (B,C,E) RNA-Seq and (D,F) RT-PCR analyses showing percent spliced in (PSI) of (B) *mbnl2* exon 5, (C-D) *aplp2* exon 7, and (E-F) *atp6v1h* exon 6 in WT and *mbnl* mutant adult zebrafish skeletal muscle. Data information: In (A) ρ is the Spearman’s rank correlation coefficient. In (A-F) homozygous *mbnl1* mutant alleles are denoted as *1^A^* and *1^B^*, *mbnl2* alleles as *2^A^*, and *mbnl3* alleles as *3^A^, 3^B^*, and *3^C^*. Representative RT-PCR gels are shown above each RT-PCR graph. Each dot represents RNA from one fish. Data are presented as mean ± SEM.

