## Appendix for "Zebrafish *mbnl* mutants model physical and molecular phenotypes of myotonic dystrophy"

### Appendix Table of Contents

**Appendix Table S1.** Zebrafish single, double, and triple homozygous *mbnl* mutants are viable to adulthood.

**Appendix Figure S1.** The introns surrounding zebrafish *mbnl1* exon 5 contain potential Mbnl protein binding sites.

**Appendix Figure S2.** Zebrafish *mbnl2* exon 5 and its surrounding introns contain potential Mbnl protein binding sites.

**Appendix Figure S3.** Zebrafish *aplp2* exon 7 and its surrounding introns contain potential Mbnl protein binding sites.

**Appendix Figure S4.** Zebrafish *atp6v1h* exon 6 and its surrounding introns contain potential Mbnl protein binding sites.

**Appendix Figure S5.** The introns surrounding zebrafish *atp2a1l* exon 23 contain potential Mbnl protein binding sites.

**Appendix Figure S6.** The introns surrounding zebrafish *ryr1b* exon 72 contain potential Mbnl protein binding sites.

**Appendix Figure S7.** The introns surrounding zebrafish *ank3b* exon 36 contain potential Mbnl protein binding sites.

| *mbnl* genotype | Cross used to generate genotype | Expected % adult offspring with genotype | Actual % adult offspring with genotype |
|---|---|---|---|
| $1^{A/A}$ | $1^{+/A}$ X $1^{+/A}$ | 25 | 28.6 (57 of 199) |
| $1^{B/B}$ | $1^{+/B}$ X $1^{+/B}$ | 25 | 25.0 (54 of 216) |
| $2^{A/A}$ | $2^{+/A}$ X $2^{+/A}$ | 25 | 28.1 (73 of 260) |
| $3^{A/A}$ | $3^{+/A}$ X $3^{+/A}$ | 25 | 15.2 (5 of 33) |
| $3^{B/B}$ | $3^{+/B}$ X $3^{+/B}$ | 25 | 23.3 (34 of 146) |
| $3^{C/C}$ | $3^{+/C}$ X $3^{+/C}$ | 25 | 29.1 (59 of 203) |
| $1^{B/B};2^{A/A}$ | $1^{+/B};2^{+/A}$ X $1^{+/B};2^{+/A}$ | 6.25 | 6.2 (13 of 209) |
| $1^{B/B};3^{C/C}$ | $1^{+/B};3^{+/C}$ X $1^{+/B};3^{+/C}$ | 6.25 | 7.9 (8 of 101) |
| $1^{B/B};3^{C/C}$ | $1^{+/B};3^{C/C}$ X $1^{+/B};3^{C/C}$ | 25 | 27.6 (8 of 29) |
| $1^{B/B};3^{C/C}$ | $1^{B/B};3^{+/C}$ X $1^{B/B};3^{+/C}$ | 25 | 27.3 (63 of 231) |
| $2^{A/A};3^{C/C}$ | $2^{+/A};3^{+/C}$ X $2^{+/A};3^{+/C}$ | 6.25 | 9.2 (13 of 142) |
| $2^{A/A};3^{C/C}$ | $2^{+/A};3^{C/C}$ X $2^{+/A};3^{C/C}$ | 25 | 25.5 (24 of 94) |
| $2^{A/A};3^{C/C}$ | $2^{A/A};3^{+/C}$ X $2^{A/A};3^{+/C}$ | 25 | 28.4 (23 of 81) |
| $1^{B/B};2^{A/A};3^{C/C}$ | $1^{+/B};2^{A/A};3^{+/C}$ X $1^{+/B};2^{A/A};3^{+/C}$ | 6.25 | 4.0 (9 of 223) |
| $1^{B/B};2^{A/A};3^{C/C}$ | $1^{+/B};2^{A/A};3^{C/C}$ X $1^{+/B};2^{A/A};3^{C/C}$ | 25 | 19.4 (55 of 284) |

**Appendix Table S1. Zebrafish single, double, and triple homozygous *mbnl* mutants are viable to adulthood.**

For each zebrafish single, double, and triple homozygous mutant *mbnl* genotype, this table lists the cross that was used to generate the genotype, the percent of adult offspring expected of the genotype according to Mendelian genetics, and the actual percent and number of adult offspring observed with the genotype.

Zebrafish *mbnl1* exon 5 / Human *MBNL1* exon 5

```
            1                                                 50
Zebrafish   gactgtatttctgcatgccttagtcctgatatccccctgtatactgcatt
Human       gccttta.ttgtgcatg.cttagtcttgttat.tcgttgtatatggcatt

            51                                               100
Zebrafish   ccaatgatttg..tttttatttgtttttttttctcacctaccc.aaaatg
Human       ccgatgatttgtttttttatttg...ttttttctcacctacccaaaaatg

            101                                              150
Zebrafish   cactgctgccccatgatgcacctctgcttgctgtttatgttaatgcgct
Human       cactgctgccccatgatgcacctctgcttgctgtttatgttaatgcgct

            151                                              200
Zebrafish   tgaaccccattggcccattgccatcatgtgctcgctgcctgctaattaag
Human       tgaaccccactggcccattgccatcatgtgctcgctgcctgctaattaag

            201                                              250
Zebrafish   ACTCAGTCGGCTGTCAAATCACTGAAGCGACCCCTCGAAGCCACCTTTGA
Human       ACTCAGTCGGCTGTCAAATCACTGAAGCGACCCCTCGAGGCAACCTTTGA

            251                                              300
Zebrafish   CCTGgtactatgacctttcaccttttgcttggcatgttgctttattgta
Human       CCTGgtactatgacctttcaccttttagcttggcatgtagctttattgta

            301                                              350
Zebrafish   g..........tgtctttaacaaacaa....aaaaactgagatataatct
Human       gatacaagtttttttttta..aatcaactttaaa.....atatata....

            351                                              400
Zebrafish   gcaagcatagtccttttgtgtttgtttaatgcctctgtt...gtgttctg
Human       ..........tcctttt...............ttctgttatagagt..tg

            401                                              450
Zebrafish   t....taccatctcgtgtg.atcaactgactgtgg.....cgagcaagaa
Human       taaagtacaa.......tgaaaaaactgagtgtggtttcctga.caa.aa

            451                                           497
Zebrafish   ttgttag.cagacgccgccgcaatc...............agatcat
Human       ttagtagaaaga.....ctataatctaagtacatagatggatatcat
```

**Appendix Figure S1. The introns surrounding zebrafish *mbnl1* exon 5 contain potential Mbnl protein binding sites.**

Zebrafish *mbnl1* exon5 (bold upper case letters) and its surrounding intronic sequences (lower case letters) were aligned with orthologous human sequences using the EMBOSS Water program. Potential YGCY (Y=pyrimidine) Mbnl protein binding sites are indicated in red.

Zebrafish *mbnl2* exon 5 / Human *MBNL2* exon 5

```
             1                                                  50
Zebrafish  ttcgtttatgtgcatgcctattgttttgtatgtggtatactgcattcaga
Human      tttgttgttgtgcatgcctagtgttttgtacgttgtatattgcattcaga

           51                                                 100
Zebrafish  ttatttttgttttgttttttttccttctcacctatccacaatgcaatgc
Human      ata.............tttttttccttctcacctatccacaatgcaatgc

           101                                                150
Zebrafish  tgtccccatgatgcacctctgcttgctg.ttacctgtatgttaatacgct
Human      cgttcccatgatgcacctctgcttgctgtttacctgtatgttaattcgct

           151                                                200
Zebrafish  tgaatcccattggcccactgccatcatgtgctcgctgcctgctattaaaa
Human      tgaatcccattggcccactgccatcatgtgctcgctgcctgttattaaaa

           201                                                250
Zebrafish  gACTCAGTCGACTGCCAAAGCAATGAAGCGACCCCTCGAGGCATCTGTAG
Human      gACTCAGTCGACTGCCAAAGCAATGAAGCGACCTCTCGAAGCAACTGTAG

           251                                                300
Zebrafish  ATCTGgtactatgacctttcacctttttgcttgcatgtagctttttaat
Human      ACCTGgtactttgacctttcacctttcgctttgcatgtagctttt.....

           301                                                350
Zebrafish  gtactgtagcagaac..tcacagaaaaaaaaagtgatttaatttggcttc
Human      ......cag.agaaccatctgag................atttg...ta

           351                                                400
Zebrafish  ttgctt.........ttgatacaaaca..................gtaa
Human      ttacttgtaaaatactggttgcaaacaatcattattattaatttatggaa

           401                                                450
Zebrafish  aaagtcaccaa....gtg.............tacttgtatgtagaactta
Human      aaaat..ccaataaggtgtttagccatctcattctcatatgta.......

           451                                        494
Zebrafish  ccatcagtcgtaattcttgaagcctagtaaacgttaagtgcctt
Human      ..atctgt.gaaataaattaatcctaacaaattttaa.taccttt
```

**Appendix Figure S2. Zebrafish *mbnl2* exon 5 and its surrounding introns contain potential Mbnl protein binding sites.**

Zebrafish *mbnl2* exon 5 (bold upper case letters) and its surrounding intronic sequences (lower case letters) were aligned with orthologous human sequences using the EMBOSS Water program. Potential YGCY (Y=pyrimidine) Mbnl protein binding sites are indicated in red.

Zebrafish *aplp2* exon 7 / Human *APLP2* exon 7

```
           1                                                 50
Zebrafish  taaa.....atac..........ag.aaatagt.ctcagc..........
Human      taaatgtctctactctgtggggaagcaacttgtcctcagcaaggctggcc

           51                                               100
Zebrafish  ........agctt.......ccggttatgcagactttcatatttttttgta
Human      tgagctggagctttcggccaccgggcctccaggct...............

           101                                              150
Zebrafish  acaaagatcttcaagtttaatatctatggagaaagtgtttacaaaaactt
Human      .......ccgtccag.......tctcagg.................cctc

           151                                              200
Zebrafish  ctgca........cccag......agtgtctgttgcgctatatattagaa
Human      ccccagcccatcccccagctcgcca..gcctgtagc.....at.tttgaa

           201                                              250
Zebrafish  ggtattgg.......tgccccgtgtggctg..gaggctctaactgcgcct
Human      .gcatttgacgtcactgcctc.tgt.cctgctgacactctgac.......

           251                                              300
Zebrafish  cctcctgtgtt......agCTGTGTGTTCTCTGGAGGCGGAGACCGGCCC
Human      ...cat...tttcacacagCTGTCTGCTCCCAGGAGGCGATGACGGGGCC

           301                                              350
Zebrafish  CTGCCGCGCCTCTATGCCTCGCTGGCACTTCGAC..ATGCAGCAGAGGAA
Human      CTGCCGGGCCGTGATGCCTCGTTGGTACTTCGACCTCTCCA..AGGGAAA

           351                                              400
Zebrafish  GTGTGTGCGCTTCATCTACGGAGGCTGCGCCGGCAACCGCAACAATTTCG
Human      GTGCGTGCGCTTTATATATGGTGGCTGCGGCGGCAACAGGAACAATTTTG

           401                                              450
Zebrafish  ACTCAGAGGAGTACTGCATGGTCGTGTG.CAAGCGCCTGAgtaagtcac.
Human      AGTCTGAGGATTATTGTATGGCTGTGTGTAAAGCG.ATGAgtaagtc.ct

           451                                              500
Zebrafish  .actcgc..........tactgaagactccgccccttctg..ctccgcct
Human      gcctcgcgctggtcccgtgcggcag.caccgtcctgtctggcgtccgtct

           501                                              550
Zebrafish  tcct.............ctgc.tccgt........cct...........t
Human      ccctgccgtcttcgtggctgcatctgtgtggtgtccctgcccactcgggt

           551                                              600
Zebrafish  ttttgct.....ccgccttcctctgctc....tgccctct.....tatgc
Human      gtttgctgtcggtcgtcttcccctcatctttgtgctttctagatctaggc

           601                                    641
Zebrafish  tctgc.ctccttctgctctgtcacagtgctaa.ctctgttc
Human      tttgctctcctgccg.......gcagtggtaagctcagttc
```

**Appendix Figure S3. Zebrafish *aplp2* exon 7 and its surrounding introns contain potential Mbnl protein binding sites.**

Zebrafish *aplp2* exon 7 (bold upper case letters) and its surrounding intronic sequences (lower case letters) were aligned with orthologous human sequences using the EMBOSS Water program. Potential YGCY (Y=pyrimidine) Mbnl protein binding sites are indicated in red.

Zebrafish *atp6v1h* exon 6 / Human *ATP6V1H* exon 6

```
             1                                                  50
Zebrafish    agactgaaa.........taaattcaaggcca..ctctcgcacagattt
Human        agaatgtaacgatatttagaacattttatgtcatcccctttgc.....ttt

             51                                                100
Zebrafish    ttagacttttgcctg.tcaagaacaaactgtctctctcatgcgggagtca
Human        tta...tttagtaggattaggaataaa........attataggggaaaaa

             101                                               150
Zebrafish    tctgtcttcctgccaccattgtccaa.....gcacacg........gctc
Human        t...tatt.......acatag...aattaggggaaaagtttgctttagctc

             151                                               200
Zebrafish    acttttt....gct.................tcaaacgaggccca.....a
Human        ccttttgtaagctgatacattaggataaaatcaaa......acaaaacta

             201                                               250
Zebrafish    aatcaccgtttttttctttctatctttttttttgttgttgt..tagAAAC
Human        aataatgagttatttatttc...................gtaatagAAAC

             251                                               300
Zebrafish    TACATGGTACAGGTGCTGAACCTGGAACAGGGACCATCTCCCCCAGTGAA
Human        TGCGTGGTAGCGGTGTTGCTGTTGAAACAGGAACAGTCTCTTCAAGTGAT

             301                                               350
Zebrafish    gtgagtat.....gtcacagatgaaggaggcagatgtgcagctgtgtgaa
Human        gtaagtatattccttcactg.......................gcctaaa

             351                                               400
Zebrafish    cgagcttccgtcccc...........tttcagaggctgag..atcttcac
Human        ........catccccaaaataaattattt.......tgagtcatataca.

             401                                               450
Zebrafish    ccctctctt....ttttgagatgcagcacatgctgttt.cttgtccactg
Human        ......cttaaaattttg...........aaggtgtttactag.......

             451                                               500
Zebrafish    cttttcccc.........cccat..tcatatccact...gtgaaaaataa
Human        ...ttccccaactataaatccatggtcacctccagttaggttaa......

             501                                               550
Zebrafish    gcatttttattacaaa..................ggtc............
Human        ....ttgtattataaactatatataaatattgatggtctgtggagatcat

             551                      576
Zebrafish    ctttcctct.............cttg
Human        cttttctgtggtagacataagacttg
```

**Appendix Figure S4. Zebrafish *atp6v1h* exon 6 and its surrounding introns contain potential Mbnl protein binding sites.**

Zebrafish *atp6v1h* exon 6 (bold upper case letters) and its surrounding intronic sequences (lower case letters) were aligned with orthologous human sequences using the EMBOSS Water program. Potential YGCY (Y=pyrimidine) Mbnl protein binding sites are indicated in red.

Zebrafish *atp2a1l* exon 23 / Human *ATP2A1* exon 22

```
           1                                                  50
Zebrafish  ~~~~~~~~~~~~~~~~~~~ttctaaaca..tgaaccttagtgtatgtttt
Human      ccgccccgccccgccccgtactttgcaggtg.........gtaagttt.

           51                                                100
Zebrafish  atataatgttttcctttttaatatacagtatattaattgca....ctttt
Human      .............ct..........cag.....ccctggcaggacctgtg

           101                                               150
Zebrafish  t........ttcctcat...............acaact............
Human      tccgccccgttccccctgcgcctgcaggggccacatctccggggcagccc

           151                                               200
Zebrafish  ...tgtttcaactaaaaagagtgtttttaatcag.......atctaattc
Human      cactgcctc....................ctcagcccccacagc....cc

           201                                               250
Zebrafish  ctata.ccccttttcttatctttctca.ctcatttaattgatttcccact
Human      ctatagccccca tgc..........cacctccctgccttga........t

           251                                               300
Zebrafish  aaccaa....ctatattcatttctag~~~~~~~~TCTAAACAGTTCTCAC
Human      aac..agtgcctcttgtcctctctggccatagGAT...AACTGTTC.CCC

           301                                               350
Zebrafish  CTCC.......CAGATTTCAAGTAACAGAAGCAAGgtatacaacacccca
Human      CTCCTCCATCTCTGAGCCCGTGTCAC.......AGgtat....cacccc.

           351                                               400
Zebrafish  gctgctt..............ttctgtg..tctg..............tc
Human      .cttcttgccctcagcccagctgctgtgcccctgccacccgcgccccctc

           401                                               450
Zebrafish  tg.....tacatgtc...tac.aggtttcaatcctttg...........c
Human      agccccttgcgcgtcgcatccaagg..tca...cttgtgctcgcagctcc

           451                                               500
Zebrafish  atgtttagtggttag..attgatgttgct....tt.........gtgg..
Human      acctggagccgtt.gccactgctgctgctgcgcttccagtcagggtgggc

           501                                               550
Zebrafish  ........tcttaaaccac......tctgttttagct...atgcatgaa
Human      cgctggcctc.....ccactgggcgtcag..tttggctcccaggc.....


           551                                               600
Zebrafish  gcgcttgtgaaagttatacttaattatacttcttaatggcaagagcttta
Human      ...cctgggcag~~~~~~~~~~~~~~~~~~~~~~~~~~~~~~~~~~~~~~

           601              618
Zebrafish  tctagttcataatttgca
Human      ~~~~~~~~~~~~~~~~~~
```

**Appendix Figure S5. The introns surrounding zebrafish *atp2a1l* exon 23 contain potential Mbnl protein binding sites.**
Zebrafish *atp2a1l* exon 23 (bold upper case letters) and its surrounding intronic sequences (lower case letters) were aligned with orthologous human sequences using the EMBOSS Needle program. Potential YGCY (Y=pyrimidine) Mbnl protein binding sites are indicated in red.

Zebrafish *ryr1b* exon 72 / Human *RYR1* exon 70

```
           1                                                 50
Zebrafish  actccatttgt......tcagacaattgctgacc............tcat
Human      actgcccttctcaggtctcagagaa....cgaccccccaccccgagccaa

           51                                               100
Zebrafish  gtcctgttgaggac.....agacactgtttctgctctcaca.ggcggtt.
Human      ggcctggaaatgcccagctagagaat............acatggcgggtg

           101                                              150
Zebrafish  ...tatagg.........gctg.....ttgatgtttttatttatttattt
Human      gggcagaggaggtggggtgctggcaacttggag.................

           151                                              200
Zebrafish  atttatttgtcctg....tctgtgagggtgtaagttttaacagtgc.tgt
Human      .....ttgggcctgggcttctctgcggggct..ggggtaacccttcttgt

           201                                              250
Zebrafish  ctctctctg.tgtcctgtcttggtgtggattcggggcctgtagGCCGGAG
Human      ctctgtctgcggtcc......ggtg............aagcagGCGGGAG

           251                                              300
Zebrafish  ATGGAGAGgtcagaccgtgtttttgcacgataaaaaaaagagctgtg..c
Human      ATATACAGgtcagccc.........cac............atctgggacc

           301                                              350
Zebrafish  tgat.cgcatggcgcagaatgttgggtttt....tcatcaaactccca..
Human      t..tccgcatgtc......tcttggctaatgccctctt....cccccagc

           351                                              400
Zebrafish  .....cacacactggcc.......aa....tgct..........cacac
Human      ctctgcacgccccccgcctcgagaaaacccctgcttctgttccccaccccc

           401                                              450
Zebrafish  atcatacacacaaaagtccaa..........accctcattcactgccagt
Human      gtcctcccc......tcccagcccccacccatcctccttcccttcc...

           451                                              500
Zebrafish  gtgtgtgaggtctgttgttgcct................cgtt.tgatga
Human      ................cttccctccccaccacccccatccgttcccatcc

           501                  521
Zebrafish  aagccctcagccatagacttt
Human      caaccctcag.......cttt
```

**Appendix Figure S6. The introns surrounding zebrafish *ryr1b* exon 72 contain potential Mbnl protein binding sites.**

Zebrafish *ryr1b* exon 72 (bold upper case letters) and its surrounding intronic sequences (lower case letters) were aligned with orthologous human sequences using the EMBOSS Water program. Potential YGCY (Y=pyrimidine) Mbnl protein binding sites are indicated in red.

Zebrafish *ank3b* exon 36 / Human *ANK3* exon 35

```
           1                                                  50
Zebrafish  ggtgcacag.gcttgtgctttttttcagcagcttctgtggttgcggcttt
Human      gatgtacagttcttgatatttttt...............gtt..ggcttt

           51                                                100
Zebrafish  tttcttttcatttg..ttgccctttatgttcct...tca.ctgtggctt
Human      tt..ttaaaatttgcttgc..........tttctaaatcagct.tgcatt

           101                                               150
Zebrafish  tgcggt...aacacaatcgcctttgttcacatcctgatgtt.tcttcccc
Human      tgtgctccctactcagt..cctt..ttcatatcctgatattctttt...c

           151                                               200
Zebrafish  cctgctcttcattacgttatcccctctgtttctgcaatgcatcttgggaa
Human      cctgctgtccattataatattcc..ttgtctctgcaatgcatctt.ggaa

           201                                               250
Zebrafish  ccaaaagGATGTGGATTCAGATCCCGAGGAAGAGgtaatagtgctgtccc
Human      ccaaaagGAGACAGAGTCAGATCAAGATGATGAGgtaata..........

           251                                               300
Zebrafish  ct...catttccatgtggtgtgctagttgctttagctct..........g
Human      .taaacacttc................tgcttttattctatcagaagtag

           301                                               350
Zebrafish  gattg..attg..tttgt...........tttaaggcggctttaacatgag
Human      aatagaaattgaatctgtatgacagaagttgcagagggc..taaagtgtg

           351                                               400
Zebrafish  tacacatacag..aacacatacagcatgctgctgcaattttctg....ca
Human      ..........ggaaacatataaattat........atttttcaggtctta

           401                                               450
Zebrafish  agttg..............attccttt.........ttcactcatccatg
Human      agtaggtaaaagggtatatatt..tttcttaaatggttcactc.ttcttc

           451                                   490
Zebrafish  cgcaagtattgttgcctggactgtcgag.acttaaa.ata
Human      cacacgt............acttaagagtacttaaatata
```

**Appendix Figure S7. The introns surrounding zebrafish *ank3b* exon 36 contain potential Mbnl protein binding sites.**

Zebrafish *ank3b* exon 36 (bold upper case letters) and its surrounding intronic sequences (lower case letters) were aligned with orthologous human sequences using the EMBOSS Water program. Potential YGCY (Y=pyrimidine) Mbnl protein binding sites are indicated in red.
